# Extending and improving metagenomic taxonomic profiling with uncharacterized species with MetaPhlAn 4

**DOI:** 10.1101/2022.08.22.504593

**Authors:** Aitor Blanco-Miguez, Francesco Beghini, Fabio Cumbo, Lauren J. McIver, Kelsey N. Thompson, Moreno Zolfo, Paolo Manghi, Leonard Dubois, Kun D. Huang, Andrew Maltez Thomas, Gianmarco Piccinno, Elisa Piperni, Michal Punčochář, Mireia Valles-Colomer, Adrian Tett, Francesca Giordano, Richard Davies, Jonathan Wolf, Sarah E. Berry, Tim D. Spector, Eric A. Franzosa, Edoardo Pasolli, Francesco Asnicar, Curtis Huttenhower, Nicola Segata

## Abstract

Metagenomic assembly enables novel organism discovery from microbial communities, but from most metagenomes it can only capture few abundant organisms. Here, we present a method - MetaPhlAn 4 - to integrate information from both metagenome assemblies and microbial isolate genomes for improved and more comprehensive metagenomic taxonomic profiling. From a curated collection of 1.01M prokaryotic reference and metagenome-assembled genomes, we defined unique marker genes for 26,970 species-level genome bins, 4,992 of them taxonomically unidentified at the species level. MetaPhlAn 4 explains ∼20% more reads in most international human gut microbiomes and >40% in less-characterized environments such as the rumen microbiome, and proved more accurate than available alternatives on synthetic evaluations while also reliably quantifying organisms with no cultured isolates. Application of the method to >24,500 metagenomes highlighted previously undetected species to be strong biomarkers for host conditions and lifestyles in human and mice microbiomes, and showed that even previously uncharacterized species can be genetically profiled at the resolution of single microbial strains. MetaPhlAn 4 thus integrates the novelty of metagenomic assemblies with the sensitivity and fidelity of reference-based analyses, providing efficient metagenomic profiling of uncharacterized species and enabling deeper and more comprehensive microbiome biomarker detection.

## Introduction

Over the last 25 years, shotgun metagenomic sequencing ^1^ and associated computational methods have developed as robust, efficient ways to study the taxonomic composition ^2–6^ and functional potential ^4,7,8^ of complex microbial communities populating human, animal, and natural environments. Genome assembly methods developed for microbial isolates have been expanded to apply to shotgun metagenomes, but while they excel in identifying new organisms from communities, their sensitivity is often limited by such environments’ complexity ^9^. Reference-based computational approaches complement assembly by relying on annotated reference sequence information to accurately identify and quantify the known taxa and genes present in a microbiome by homology instead ^4–7^. This set of methods enabled deep exploration of human microbiomes and the discovery of microbial associations with multiple health conditions ^10–18^ and dietary patterns ^19–23^, as well as the characterization of the evolution and transmission of microbial species and strains ^24–29^. However, reference-based methods can only detect well-characterized and cataloged microbial species included in available reference databases, which typically only represent a fraction of the community members across environments, thus limiting the interpretation of shotgun metagenomes ^30^.

Conversely, *de novo* metagenomic assembly to reconstruct draft genes and genomes - called metagenome-assembled genomes (MAGs) - has advanced to the point of very high specificity (albeit often low sensitivity) for recovery directly from metagenomes ^31–35^. This allows recovering of microbial sequences that have not yet been isolated or characterized and thus absent from reference databases ^36^. As metagenomic assembly and binning have improved dramatically in the last few years ^31–35^, large-scale MAG catalogs have been compiled and comprise a vast amount of unknown and uncultivated microbial species populating diverse environments ^37–46^. However, such metagenomic assembly techniques are typically able to capture only a limited fraction of the organisms in complex communities due to insufficient coverage for many taxa, the presence of genetically related taxa impeding or creating spurious assemblies, and difficulties in quality control of the resulting MAGs ^9^.

To leverage the best aspects of both reference- and assembly-based metagenome profiling, we present MetaPhlAn 4, a method that exploits an integrated extended compendium of microbial genomes and MAGs to define an expanded set of species-level genome bins ^42^ (SGBs) and accurately profile their presence and abundance in metagenomes. SGBs represent both existing species (known or kSGBs) or yet-to-be-characterized species (unknown, uSGBs) defined solely based on MAGs ^42^. From a collection of 1.01M bacterial and archeal MAGs and isolate genomes spanning multiple environments, we first expanded the definition of 54,596 SGBs, and then defined SGB-specific unique marker genes (i.e. genes uniquely characterizing each SGB) for 21,978 kSGBs and 4,992 uSGBs. The resulting dataset expands the existing MetaPhlAn algorithm ^2–4^ to enable deeper and more accurate quantitative taxonomic analyses of human, host associated, and environmental microbiomes and provides new insights in a number of studies associating the microbiome with host conditions.

## Results

### Taxonomic profiling of metagenomes using combined metagenomic assemblies and isolate references

MetaPhlAn 4 expands and improves existing capabilities to perform taxonomic profiling of metagenomes by exploiting a framework in which extensive metagenomic assemblies are integrated with existing bacterial and archaeal reference genomes. These are then jointly preprocessed to allow efficient metagenome mapping against millions of unique marker genes, ultimately quantifying both isolated and metagenomically-assembled organisms in new communities. The algorithm augments that used by previous versions in four main ways: (i) the adoption of species-level genome bins (SGBs) ^42^ as primary taxonomic units, each of which groups microbial genomes and metagenome assembled genomes (MAGs) into consistent existing species and newly defined genome clusters of roughly species-level diversity; (ii) the integration of over 1M MAGs and genomes into this SGB structure to build the largest database of confident microbial reference sequences currently available; (iii) the curation of microbial taxonomic units based on the consistency of taxonomically-labeled microbial genomes and the assignment of new taxonomic labels to SGBs solely defined on MAGs; (iv) the improved procedure to extract unique marker genes out of each SGB for the MetaPhlAn reference-based mapping strategy ^2–4^. MetaPhlAn 4 thus leverages aspects of both metagenomic assembly, with its potential to uncover previously unseen taxa ^40–42,45^, and the sensitivity of reference-based profiling to provide accurate taxonomic identification and quantification.

The adoption of SGBs as the primary unit of taxonomic analysis is central to this approach ^42^. Briefly, an SGB ^42^ delineates a microbial species purely based on the clustering of whole-genome genetic distances at 5% genomic identity ^47^ and a taxonomic label can then be assigned to the SGB based on the presence (or not) of characterized genomes from isolate sequencing. This definition permits arbitrary microbial genomes to be organized in a manner not unlike amplicons into operational taxonomic units (OTUs) and matches remarkably well the expected boundaries of the existing taxonomy ^42,47,48^. Available microbial reference genomes and medium-to-high-quality MAGs are thus grouped into taxonomically well-defined species (“known” SGBs or kSGBs when a isolate genome with available taxonomy is present in the SGB) or unknown equivalent clades (uSGBs).

Following the SGB clustering approach, the database employed by MetaPhlAn 4 contains SGBs that result from the merging of species that were originally incorrectly taxonomically labeled as separate species. For example, genomes assigned in NCBI ^49^ to *Lawsonibacter asaccharolyticus* and *Clostridium phoceensis* are 98.7% identical, likely due to independent naming of members of a new species, and were merged into the SGB15154 (**Table S1**). This merging also applies to taxonomic species that are genetically difficult or impossible to distinguish (e.g. species of the *Bacillus cereus* group, genetically differentiated only by their plasmidic sequences ^50^) and are thus clustered in the same SGB. Conversely, species with subclades diverging for more than 5% genetic identity were splitted into multiple SGBs (e.g. *Prevotella copri* is represented by four different SGBs ^51^, or *Faecalibacterium prausnitzii* with SGBs representing its distinct (sub)species ^52^, **Table S1**). Finally, incorrectly or partially taxonomically classified reference genomes were detected and amended based on the detection of outlier labels resulting from misspellings or incorrect assignments by NCBI genome submitters (e.g. the *Staphylococcus epidermidis* SGB7865 is composed of 700 reference genomes, 32 of which have different or unspecified species labels in the NCBI database ^49^, **Table S1**).

To derive the database of SGBs to be profiled in MetaPhlAn 4, the isolate genome component included 236,620 bacterial and archeal genomes available in NCBI ^53^ and labeled as “reconstructed from isolate sequencing or single cells”. These were integrated with 771,528 MAGs assembled from samples collected from humans (5 distinct main human body sites, 164 distinct human cohorts), animal hosts (including 22 non-human primate species), and non-host associated environments (including soil, fresh water, and oceans, **Tables S2 and S3**). After removing reference genomes and MAGs that did not meet quality control criteria (i.e. genome completeness above 50% and contamination below 5%, see **Methods**), the catalog comprised 729,195 genomes (560,084 MAGs and 169,111 reference genomes) and was Mash ^54^ clustered into SGBs at 5% sequence similarity ^42^ for the final database of 70.9k SGBs, 47.6k of which are taxonomically unknown at the species level (uSGBs) (**Fig. 1a**). To decrease the potential rate of false positive detection of SGBs without strong support or that are extremely rare, we retained only the uSGBs containing at least 5 MAGs from distinct samples for subsequent metagenome profiling, resulting in a final catalog of 29.4k quality-controlled SGBs (see **Methods**).

**Figure 1.**
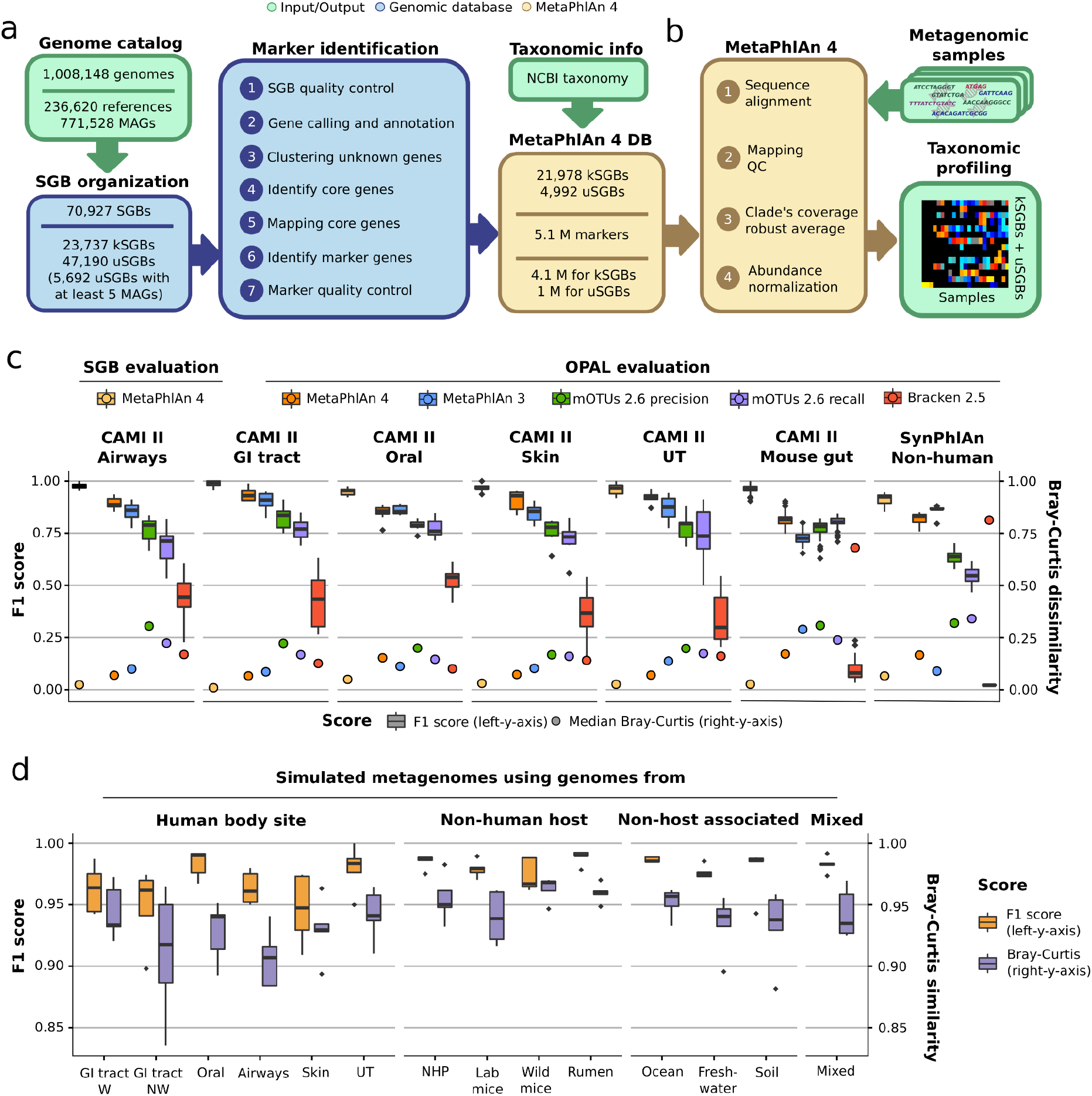
MetaPhlAn 4 improves sensitivity and specificity of metagenome taxonomic profiling by integrating reference sequences from isolate and metagenome-assembled genomes. **a**, From a collection of 1.01M bacterial and archeal reference genomes and metagenomic-assembled genomes (MAGs) spanning 70,927 species-level genome bins (SGBs), our pipeline defined 5.1M unique SGB-specific marker genes that are used by MetaPhlAn 4 (avg. 189 ± 34 / SGB). **b**, The expanded marker database allows MetaPhlAn 4 to detect the presence and estimate the relative abundance of 26,970 SGBs, 4,992 of which are candidate species without reference sequences (uSGBs) defined by at least 5 MAGs. The profiling is performed firstly by (1) aligning the reads of input metagenomes against the markers database, then (2) discarding low-quality alignments and (3) calculating the robust average coverage of the markers in each SGB that (4) are normalized across SGBs to report the SGB relative abundances (**Methods**). **c**, To evaluate its performance in taxonomic profiling, MetaPhlAn 4 was applied to 118 synthetic metagenomes representing host-associated communities from the CAMI II taxonomic profiling challenge ^57^ and the SynPhlAn-nonhuman dataset, representing more diverse environments from previous evaluations ^4^. Species-level evaluation using the OPAL framework ^58^ shows that MetaPhlAn 4 is more accurate than the available alternatives in both the detection of which taxa are present (the F1 score is the harmonic mean of the precision and recall of detection) and their quantitative estimation (the Bray-Curtis beta-diversity is computed between the estimated profiles and the abundances in the gold standard). Additional evaluations performed using genomes within the SGB organization (labeled “SGB evaluation”, see **Methods**) show that MetaPhlAn 4 further improves accuracy at this more refined taxonomic level. See **Tables S4-5** for more details (GI=gastrointestinal, UT=urogenital tract). **d**, MetaPhlAn 4 was applied to 70 synthetic metagenomes modeling different host and non-host associated environments and containing, on average, 47 genomes from both kSGBs and uSGBs (see **Methods**). This evaluation directly on SGBs shows the reliability of MetaPhlAn 4 to quantify both known and unknown microbial species. Additional evaluation based on a mixture of novel MAGs from samples not considered in the building of the genomic database (Mixed evaluation) stresses its accuracy independently from the inclusion of the profiled data in the database. See **Tables S6-7** for more details (NHP=non-human primates). All data are presented as mean ± s.d.

From this SGB genome catalog, we built the pangenome of each SGB (collection of all gene families found in at least one genome in the SGB) and used them to identify species-specific marker genes for MetaPhlAn profiling. The pangenomes were built by categorizing the coding sequences of all the 729k genomes into UniRef90 clusters ^55^ when a 90% amino acid identity match was found within the UniRef database, or by *de novo* clustering all remaining sequences at 90% amino acid identity following the Uniclust90 criteria ^56^ (see **Methods**). From the resulting 50.6M UniRef90 identities and 77.7M new Uniclust90 gene families, we subsequently identified core gene families (i.e. those present in almost all genomes and MAGs of an SGB, see **Methods**) and then screened them for their species-specificity by mapping against all sequences of all SGBs (see **Methods**). This procedure resulted in 5.1M total unique marker genes spanning 26,970 high quality SGBs, with an average of 189 ± 34 unique marker genes per SGB. MetaPhlAn 4 taxonomic profiling uses these markers to detect the presence of an SGB (known or unknown) in new metagenomes based on the detection via read mapping of a sufficient fraction of SGB-specific marker genes (default 20%) and quantifies their relative abundance based on within-sample-normalized average coverage estimations (see **Methods, Fig. 1b**).

### MetaPhlAn 4 improves the performance of metagenomic taxonomic profiling

To evaluate the taxonomic profiling performance of MetaPhlAn 4, we first assessed its ability to profile well-characterized species (i.e. those belonging to kSGBs) in comparison with available methods by using 123 synthetic metagenomes (4.1B total reads). Most of these synthetic samples (118) are from the CAMI II taxonomic profiling challenge ^57^ representing host-associated communities, while the other 5 are additional non-human synthetic metagenomes (derived from SynPhlAn, see **Methods**) representing more diverse environments than in previous evaluations ^4^.

Through the OPAL benchmarking framework ^58^, we evaluated MetaPhlAn 4 in comparison with MetaPhlAn 3 ^4^, mOTUs 2.6 ^6^ (latest database available as for March 2021) and Bracken 2.5 ^5^ (with the database built using the April 2019 RefSeq release ^59^). MetaPhlAn 4 outperformed the other tools when assessing the F1 score (**Fig. 1c**) computed based on the common reference NCBI taxonomy. This was true despite the fact that OPAL does not consider SGB-defined species groups (i.e. single species incorrectly taxonomically labeled as separated species and included in the same SGB), thus penalizing MetaPhlAn 4 profiling that cannot match the corresponding labels; the new version still achieved a higher number of species correctly detected compared with MetaPhlAn 3 across all simulations (avg 83.02 ± 47.38 and 70.62 ± 35.85 true positives, respectively) while maintaining a low number of false positives (avg 11.61 ± 8.4 and 8.69 ± 4.89, respectively, **Supplementary Fig. 1, Table S4**). Most of the false positives (84.6%) were due to the new labels of SGB-defined species groups (e.g. the *Marinilactibacillus* sp. 15R, present in almost all the CAMI II oral metagenomes, belongs to the *Marinilactibacillus piezotolerans* SGB7875 species group) and are thus also not strictly false positives. The improvement in recall may be explained by the expanded catalog of reference genomes included in MetaPhlAn 4 (169.1k genomes spanning 31.9k species in comparison with 99.2k genomes from 13.5k species in MetaPhlAn 3).

We then evaluated the relative abundance quantification performance of MetaPhlAn 4 using Bray-Curtis dissimilarity with respect to synthetic reference community compositions. MetaPhlAn 4 outperformed all alternative methods in all environments (avg. Bray-Curtis distance to the gold standard abundances 0.14 ± 0.07), including the previous MetaPhlAn version 3 (avg 0.21 ± 0.12, **Table S5, Fig. 1c**). The quality of the marker set is likely the driving factor of this improvement, a consequence of the phylogenetic consistency of the SGBs that ensures that identically-labeled taxa are genomically consistent. This avoids hard-to-detect taxonomic mislabellings in the original, manually-assigned taxonomic labels and allowed us to obtain a larger (avg 189 ± 34 per SGB as compared to 84 ± 47 per species in MetaPhlAn 3), reliable (**Supplementary Fig. 2**), and more unique set of marker genes (99.3% of the markers in comparison with 72.7% in MetaPhlAn 3).

Apropos, since these evaluations were not able to account for modifications of species taxonomy that avoid these issues, we then evaluated MetaPhlAn 4 on the same synthetic metagenomes, but using SGB-based taxonomy (see **Methods**). By considering as the gold standard label for each genome in the synthetic community the SGBs it belongs to, MetaPhlAn 4 achieved high accuracies when assessing both the F1 score (avg 0.96 ± 0.024) and the Bray-Curtis dissimilarity (avg 0.034 ± 0.022, **Fig. 1c**).

Finally, we assessed the performance of MetaPhlAn 4 to specifically detect uSGBs representing clades without taxonomically characterized isolates. We constructed 65 synthetic metagenomes simulating microbiomes from 12 different human body sites, animal hosts, and non-host associated environments, using both kSGBs and uSGBs that were found and reconstructed in real metagenomes in each of the environments via metagenomic assembly (see **Methods**). We also built 5 additional synthetic metagenomes using a mixture of MAGs and reference genomes from samples not included in our original genomic database (see **Methods**). MetaPhlAn 4 showed accuracies in the detection and quantification of uSGBs (avg. F1 score 0.97 ± 0.02, **Fig. 1d, Supplementary Fig. 3**) that were on par with those of known species (kSGBs, avg. F1 score 0.96 ± 0.024, **Fig. 1c**). Both the F1 score and the Bray-Curtis similarity to the gold standard were consistent across all the different environments assessed. Synthetic samples based on MAGs not available at the time when the MetaPhlAn 4 database was built yielded similar results (avg. F1 score 0.98 ± 0.006, **Fig 1d, Tables S6-7**). Altogether, MetaPhlAn 4 outperformed the other available tools on synthetic data and further provided quantification of yet-to-be-characterized species, while maintaining high accuracy for taxonomically well-defined species.

### MetaPhlAn 4 expands the profiled fraction of metagenomes

The MetaPhlAn 4 database expands the number of quantifiable known microbial species (18.4k more species than in MetaPhlAn 3), refines the resolution of many species described by kSGBs (21,978 kSGBs, with avg. 1.15 kSGBs per species), and includes 4,992 yet-to-be-characterized microbial species (uSGBs). We assessed its resulting increased ability to explain a larger fraction of the reads in a metagenome by profiling a total of 24.5k metagenomic samples (145 distinct studies, **Table S8**) from different human, animal, and non-host associated environments (**Fig. 2a, Supplementary Fig. 4**). We further divided the 19.5k human metagenomes based on the body site of origin and the lifestyle (i.e. Westernized or non-Westernized) of the donor (for a full description of Westernization, see **Methods**).

**Figure 2.**
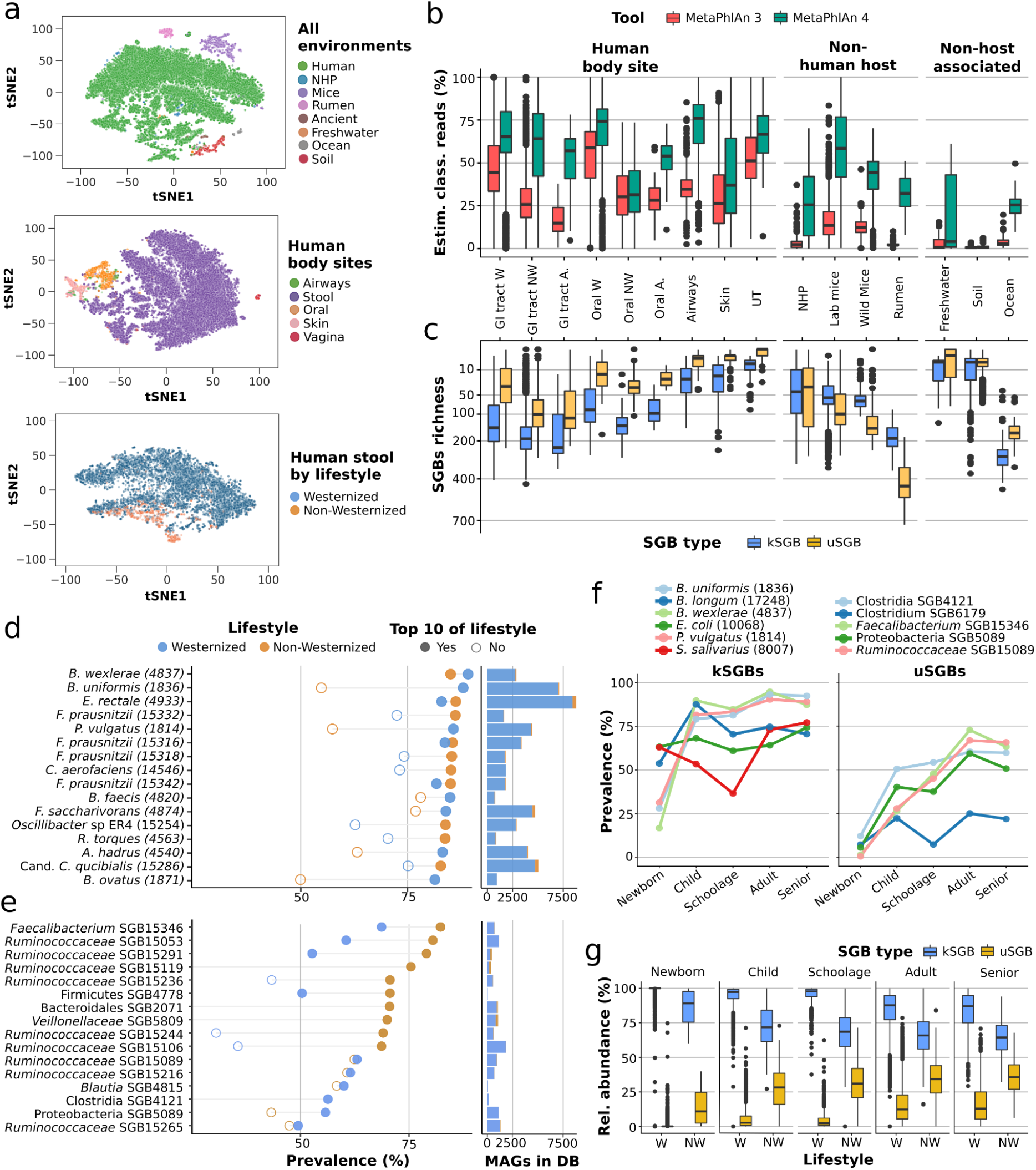
MetaPhlAn 4 expands observable microbial diversity, primarily by quantifying yet-to-be-characterized species (uSGBs). **a**, We applied MetaPhlAn 4 to a total of 24.5k metagenomic samples from different human body sites and lifestyles, animals, and non-host associated environments (**Supplementary Fig. 6a, Table S8**). **b**, The expanded genomic database of MetaPhlAn 4 significantly increases the estimated fraction of classified reads in comparison with the previous MetaPhlAn version across habitat types (GI=gastrointestinal, UT=urogenital tract, NHP=non-human primates, W=Westernized, NW=non-Westernized, A=ancient). **c**, MetaPhlAn 4 detects on average 48 unknown bacterial species (uSGBs) per human gut microbiome, and reaches up to more than 700 in other non-human environments (e.g. individual rumen samples). **d**, The most prevalent microbial species in the gastrointestinal tract of Westernized populations are known species (kSGBs). The 10 most prevalent kSGBs in Westernized and non-Westernized lifestyles (bold dots) are shown ordered by their highest prevalence and reported together with the number of MAGs assembled from human gut metagenomes in the MetaPhlAn genome catalog. Species names are shown together with their SGB ID between brackets. **e**, The most prevalent SGBs in non-Westernized populations belong to yet-to-be-cultivated and named species. The 10 most prevalent uSGBs of each lifestyle (bold dots) are shown ordered by their highest prevalence. **f**, In Westernized populations, the most prevalent kSGBs and uSGBs vary across age categories. The 2 most prevalent SGBs for each age category are shown. **g**, The fraction of uSGBs relative to kSGB increases after infancy (W=Westernized, NW=non-Westernized).

In the resulting taxonomic profiles, MetaPhlAn 4 detected 11,132 SGBs present in at least 1% of the samples of one of the environments, 3,527 of which (31.68%) were taxonomically unknown at the species level (uSGBs). The new profiles explained a much larger fraction of the reads in the metagenomic samples compared to the previous version across all environments (**Fig. 2b**). Within the human body sites, the improvement was high for the airways (avg. 1.95 fold increase of explainable reads), and substantially higher improvements were reached for samples from e.g. the gut microbiomes of non-human mammals, ranging from avg. 3.26 fold increase of the wild mice to 14.15 fold increase of the rumen. For these animals, the average number of uSGBs detected surpassed that of the kSGBs (with the exception of the non-human primates, **Fig 2c, Supplementary Fig. 5)**. These increases were consistent with the number of newly considered MAGs that defined new uSGBs from non-human microbiomes (90,606 MAGs defining 1,287 uSGBs).

Environmental ecosystems had metagenomes that were generally less explained by the taxa considered in MetaPhlAn 4, with soil in particular remaining poorly characterized due its remarkable microbial variability and the lack of systematic large metagenomic efforts targeting it (only 2,495 MAGs defining 26 uSGBs in our database), while the ocean microbiome had a 6.65 fold increase, largely due to the inclusion of the Tara ocean MAGs ^60^ in the SGB database (**Fig 2c**). Overall, uSGBs were instrumental to increase the fraction of metagenomes profileable by MetaPhlAn 4 (**Fig 2b**), as they accounted for an average of 23.13 % (s.d. 17.89%) of the richness of the resulting profiles across all environments (**Fig 2c**).

### SGB profiling reveals species overlaps across environments and potential common contaminants

A key advantage of reference-based metagenomic profiling as compared to assembly is its ability to detect low-abundant and hard-to-assemble genomes ^37,39,40,42,43,45^. This allows the generation of confident ecological statistics regarding prevalent and rare taxa, which are difficult to quantify accurately in the presence of many technical non-detections in data solely from metagenome assemblies. On this dataset, MetaPhlAn 4 identified 1,657 SGBs found in at least 1% of the samples from the gut of non-Westernized human populations (550 of these being uSGBs), 331 SGBs at the same prevalence threshold in the typically low-diverse human vaginal microbiome (61 of which are uSGBs), and intermediate numbers in other environments (**Supplementary Fig. 6a**).

This confirmed that gut metagenomes retrieved from ancient samples (ranging from 5,300 to 150 years ago in the available datasets) possessed more SGBs in common with those at >1% prevalence in the gut microbiome of modern non-Westernized populations (1,039 SGBs) than of Westernized ones (748 SGBs), despite the dominance in datasets and databases of data derived from Westernized populations (∼ten times more samples, **Supplementary Fig. 6a, Table S3**). Similarly, and adopting the same prevalence threshold at 1%, the SGBs found in the gut of non-human primates (including those in captivity) overlapped more with gut samples from ancient microbiomes (879 SGBs) than with modern ones (668 SGBs), further highlighting the effect of lifestyle in shaping the human microbiome (**Supplementary Fig. 6a**). A similar environmental adaptation can be observed in the gut microbiome of laboratory mice, in which many more modern human gut SGBs were found (481 SGBs) compared to those from wild mice (53 SGBs). Twenty-eight SGBs were present at >1% prevalence in all human body sites (**Table S9**), comprising typically oral microbes that can reach the lower gastrointestinal tract, can contaminate the skin, and can colonize other mucosal sites such as the vagina, i.e. the *Haemophilus parainfluenzae* group (SGB9712), the *Streptococcus salivarius* group (SGB8007), *Veillonella parvula* (SGB6939), *Rothia mucilaginosa* (SGB16971), and *Streptococcus oralis* (SGB8130).

Species overlap across environments at the same 1% prevalence threshold can also spot potential contamination, as it is the case of the only 9 SGBs shared between the modern human gut and ocean water samples (**Table S10**). These were predominantly skin and oral microbes likely to contaminate low-biomass water samples during laboratory procedures: *Cutibacterium acnes* (SGB16955), *Staphylococcus aureus* (SGB7852), *Streptococcus thermophilus* (SGB8002), *Escherichia coli* (SGB10068), *V. parvula* (SGB6939), *Staphylococcus epidermidis* (SGB7865), *Staphylococcus hominis* (SGB7858), *Streptococcus mitis* (SGB8163), and *R. mucilaginosa* (SGB16971). Overall, the new MetaPhlAn 4 profiling highlights that microbiomes from most non-host associated environments have little overlap between themselves and the human microbiome (**Supplementary Fig. 6b**), and that, as expected, human microbiomes from different body sites have limited but relevant overlaps (**Table S9)**.

### MetaPhlAn 4 expands the panel of highly prevalent human intestinal species

We assessed the prevalence of SGBs in the gut microbiome of human individuals (**Table S11**) using 19.5k human gut metagenomes from 86 datasets, spanning different age categories, geographic locations and lifestyles (**Table S12)**. The most prevalent SGBs in Westernized populations were from known species (**Fig. 2d**), specifically *Blautia wexlerae* (SGB4837, 89.2%), the *Bacteroides uniformis* group (SGB1836, 88.1%), and *Phocaeicola vulgatus* (previously *Bacteroides vulgatus*, SGB1814, 85.8%). Four distinct *F. prausnitzii* SGBs appeared within the top 10 most prevalent species, and three of them had quite distinct prevalence in both lifestyles (**Fig. 2d**), highlighting the ability of SGB profiling to increase the resolution of species that are particularly genetically divergent ^52^. *Cibionibacter quicibialis* ^42^, as well as several other species of interest considered kSGBs because they have a sequenced representative even though they remain largely uncharacterized (e.g. *Oscillibacter sp*. ER4) were also found at high prevalence (**Fig. 2d**).

While most uSGBs had lower prevalence in this population, 4 uSGBs from the *Ruminococcaceae* family exceeded 75% prevalence, and many of them were substantially more prevalent in non-Westernized compared to Westernized populations (**Fig. 2e**). The species with the highest prevalence in each specific age category displayed variable prevalence in the other age groups (**Fig. 2f, Supplementary Fig. 7, Table S11**), and uSGBs tended to be particularly common in childhood, which may be under-studied relative to infancy and adulthood (**Fig. 2g, Table S13**). Overall, the newly established SGBs prevalence across population and lifestyles (**Table S11**) expands both the size and detail of that established by prior metagenomic studies.

### Biomarkers of diet in mice are dominated by uSGBs

MetaPhlAn 4 integrates 22,718 MAGs assembled from 1,906 mouse gut metagenomes (both research laboratory mice and wild mice) and defines 540 uSGBs, allowing greater resolution in profiling the murine gut. When applied to a heterogeneous public dataset of 184 mouse gut microbiomes spanning 8 genetic backgrounds and 6 different vendors (**Table S14**) ^61^, MetaPhlAn 4 detected 632 different SGBs, 60.8% of which were uSGBs (**Fig. 3a, Supplementary Fig. 8a**). In contrast, only 108 total species were detected by MetaPhlAn 3 from the same samples. Interestingly, of the 43 SGBs present in more than 75% of the samples, most are uSGBs; the 12 kSGBs themselves represent poorly-characterized species such as *Lachnospiraceae* bacterium 28_4 (SGB7272), *Dorea* sp. 5_2 (SGB7275) and *Oscillibacter* sp. 1_3 (SGB7266), which were also the only ones detectable by MetaPhlAn 3. The poor mappability of many mouse microbiomes against isolate genomes is also reflected at taxonomic levels higher than species, as more than half of the families (i.e. FGBs defined similarly to SGBs but spanning up to 30% nucleotide divergence, see **Methods**) present in more than 20% of the samples are still uncharacterized (uFGBs, **Fig. 3b, Supplementary Fig. 8b**).

**Figure 3.**
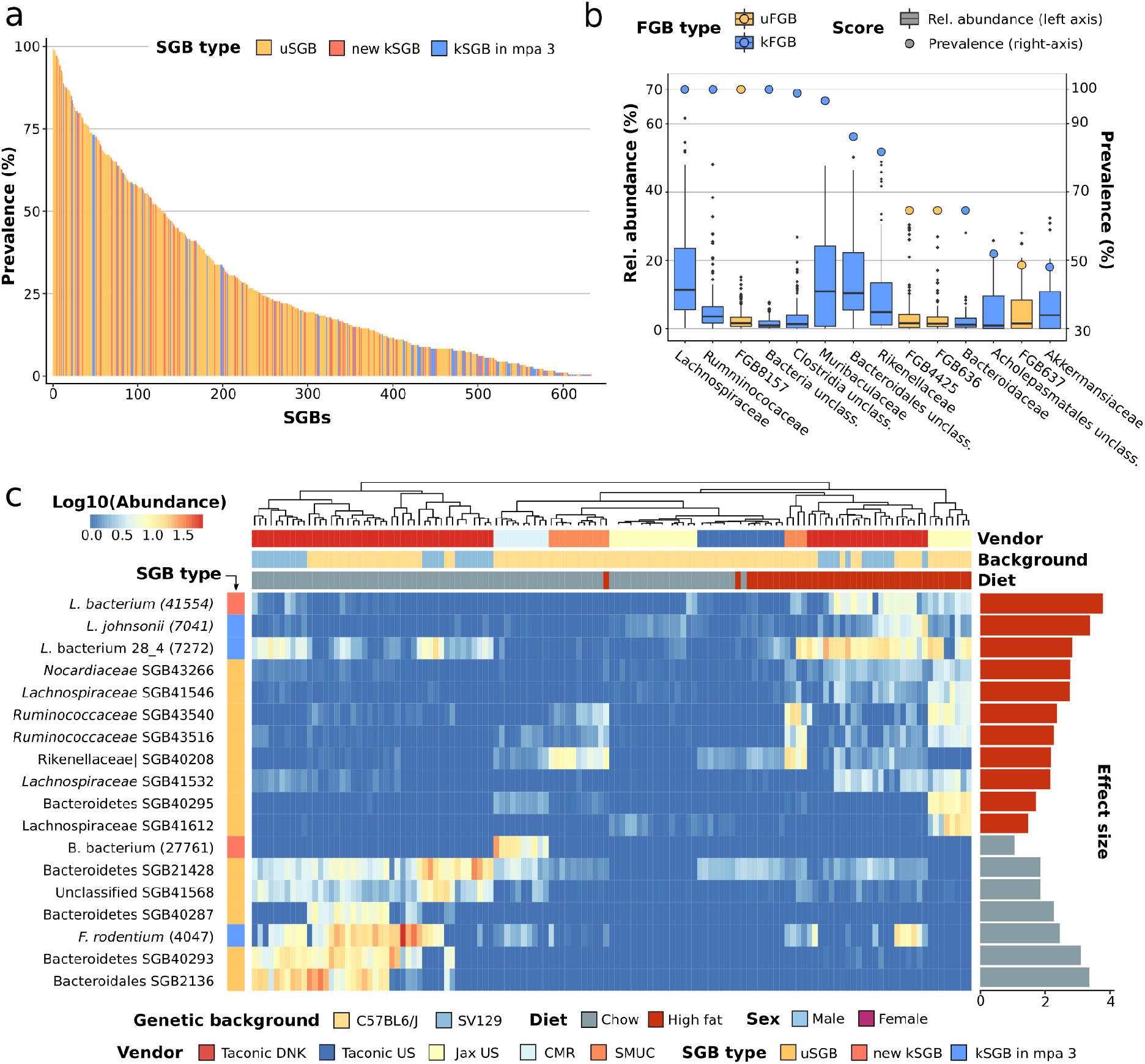
MetaPhlAn 4 enables accurate metagenomic profiling of mouse microbiomes containing few cultured isolate taxa. **a**, MetaPhlAn 4 taxonomic profiling of a cohort of 181 mouse gut microbiome samples, spanning 8 genetic backgrounds and 6 different vendors ^61^ revealed that the majority of detected microbial taxa are uncharacterized SGBs (uSGBs) that do not contain a sequenced isolate representative. **b**, Some of the most prevalent families in the mouse gut microbiome are still unclassified at the family level (uFGBs). FGBs detected in at least 20% of the samples (circles and right-side *y* axis) and with a median relative abundance above 1% (box plots and left-side *y* axis) are shown. **c**, Random effects models applied to the MetaPhlAn 4 profiles revealed that most of the high- and low-fat diet microbial biomarkers are uncharacterized species (FDR<0.2). Log10-transformed relative abundances of the microbial biomarkers are represented in the heatmap and their effect size (linear model beta coefficient) in the bar plots. For kSGBs, species names are shown together with their SGB ID between brackets. SMUC=Southern Medical University in China, CMR=Craniofacial Mutant Resource at the Jackson Laboratory (Jax).

To test the relevance of uSGBs in the context of typical mouse microbiome studies, we recapitulated prior statistical tests to identify taxonomic biomarkers of high-fat (HF) versus normal chow diets across host genetic backgrounds and vendors ^61^. Applying linear mixed models on the MetaPhlAn 4 taxonomic profiles and controlling for sex, age, genetic background, and vendor (see **Methods, Table S15)**, we identified 18 SGB biomarkers at FDR < 0.2 with an average relative abundance in the associated diet >1% (**Fig. 3c**). Most of the over-abundant biomarkers of a hyper-caloric diet were uSGBs (13 uSGBs, 72% of the 18 biomarkers), in addition to 3 taxa that could be detected using MetaPhlAn 3 (*Lachnospiraceae* bacterium 28_4 SGB7272, *Lactobacillus johnsonii* SGB7041, and *Faecalibaculum rodentium* SGB4047) and 2 kSGBs representing poorly characterized species (*Lachnospiraceae* bacterium SGB41544 and Bacteroidales bacterium SGB27761). MetaPhlAn 4’s ability to profile species defined solely by MAGs (i.e. uSGBs) thus appears particularly relevant for under-characterized microbial environments in which cultivated and sequenced taxa still represent a small fraction of overall microbial diversity.

### Stronger links between the human gut microbiome, host diet, metabolism

We used MetaPhlAn 4 to extend links between the gut microbiome, diet, and host metabolism ^19–23,62^ by re-analyzing metagenomes from 1,001 deeply phenotyped individuals in the ZOE PREDICT 1 study ^22^. As in the original study, strengths of association between the microbiome and both dietary and cardiometabolic host variables were evaluated by testing the predictive power of random forest (RF) classifiers and regressors trained on the taxonomic profiles (see **Methods**). Among the 19 health and diet markers most strongly linked with the microbiome according to MetaPhlAn 3 in the original work, all but two were better-predicted when incorporating MetaPhlAn 4 taxa (new median AUC=0.74, 4.84% improvement, **Fig. 4a**). The highest improvement was found for the 10-year atherosclerotic cardiovascular disease (ASCVD) risk (0.106 higher AUC, 16.24% improvement), and the Healthy Eating Index (HEI) score ^63^ achieved the strongest association (0.072 higher AUC, 10.05% improvement, and 31% regression improvement).

**Figure 4.**
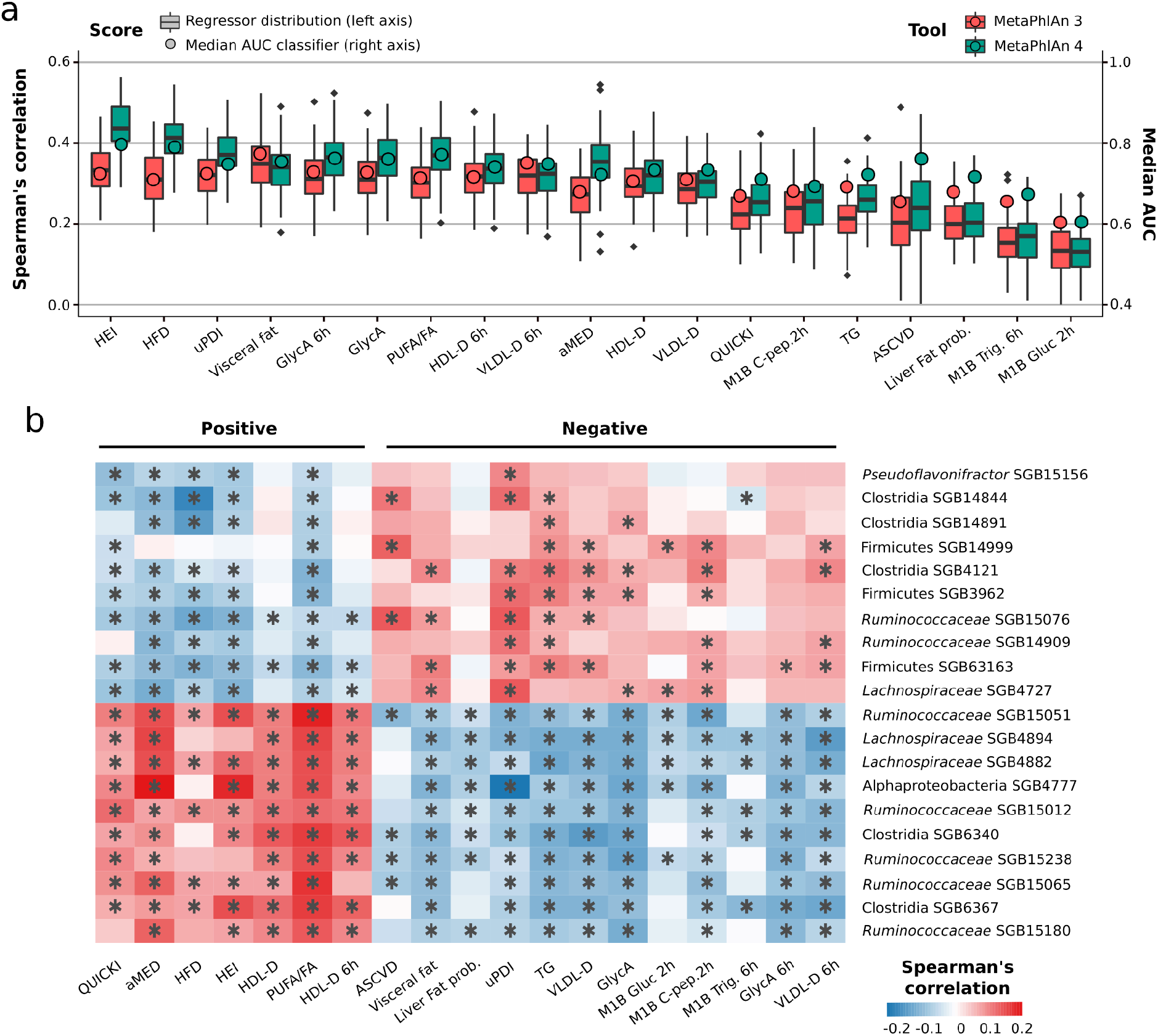
MetaPhlAn 4 reveals strong links between the unknown fraction of the human gut microbiome and host diet and cardiometabolic markers. **a**, Compared to the original results from the ZOE PREDICT 1 study based on the MetaPhlAn 3 taxonomic profiles ^22^, random forest (RF) models trained on the MetaPhlAn 4 microbiome profiles significantly improve classification (circles and right-side *y* axis) and regression (box plots and left-side *y* axis) results for a panel of 19 markers representative of nutritional and cardiometabolic health (see **Methods**). **b**, Panel of the 20 unknown microbial species (uSGBs) showing the strongest overall correlations with the positive (bottom-half list) and negative (top-half list) dietary and cardiometabolic health markers, respectively (∗ = FDR<0.2).

Microbiome links with dietary indices were particularly improved by considering uSGBs (**Fig. 4a**); previously, visceral fat and blood lipid levels were generally more strongly microbiome-associated than dietary indices using MetaPhlAn 3 profiles. This was substantiated by the analysis of correlation between the abundance of each uSGB with all 19 host diet, anthropometric, and physiology indices. Indeed, the strongest correlations (after accounting for age, sex, and BMI, **Fig. 4b**) mostly involved uSGBs (6 out of the 10 SGBs most associated with healthy conditions were uSGBs), and the three highest (absolute) correlations involved Alphaproteobacteria SGB4777, positively correlating with the alternate Mediterranean diet (aMED ^64^, *p*=0.21) and HEI (*p*=0.19) scores, and negatively correlating with the uPDI (*p*=-0.25).

We further compared the SGBs newly linked to diet and biometrics in the ZOE PREDICT 1 re-analysis to those associated with other health and disease conditions in our broader human gut data MetaPhlAn 4 profiles. Among the 10 uSGBs most health-associated based on the average correlation ranks with the 19 reference markers selected from the ZOE PREDICT 1 study, *Lachnospiraceae* SGB4894 emerged as a particularly relevant taxon. This uSGB was prevalent in both contemporary human cohorts (44.33% in healthy individuals) and in non-human primates (41.36% prevalence). It was also present in 60% of the metagenomes available from ancient stool samples (**Supplementary Fig. 9a**), suggesting that this taxon is an important, as-yet-uncharacterized member of the healthy human microbiome.

When comparing the relative abundances of *Lachnospiraceae* SGB4894 in case/control studies across datasets spanning 11 different human diseases (see **Methods, Table S16**), we found statistically significant associations not only with conditions directly linked with cardiometabolic health such as ASCVD (p-value=0.045) and cirrhosis (p-value=9.20e-7), but also with the inflammatory bowel diseases (IBD) (**Fig. 5a**). This included associations over three different cohorts of a higher abundance and prevalence of *Lachnospiraceae* SGB4894 with both of the main IBD subtypes, Crohn’s disease (p-values=2.50e-28, 4.67e-6, and 0.0016) and ulcerative colitis (p-values=1.85e-22, 3.89e-6, and 1.28e-8). Altogether, these results show the importance of profiling the unknown fraction of the microbiome even for relatively well-characterized environments such as the human gut, as microbial links with cardiometabolic blood metabolites, dietary patterns, and host diseases can also incorporate and shed light on newly defined uSGBs.

**Figure 5.**
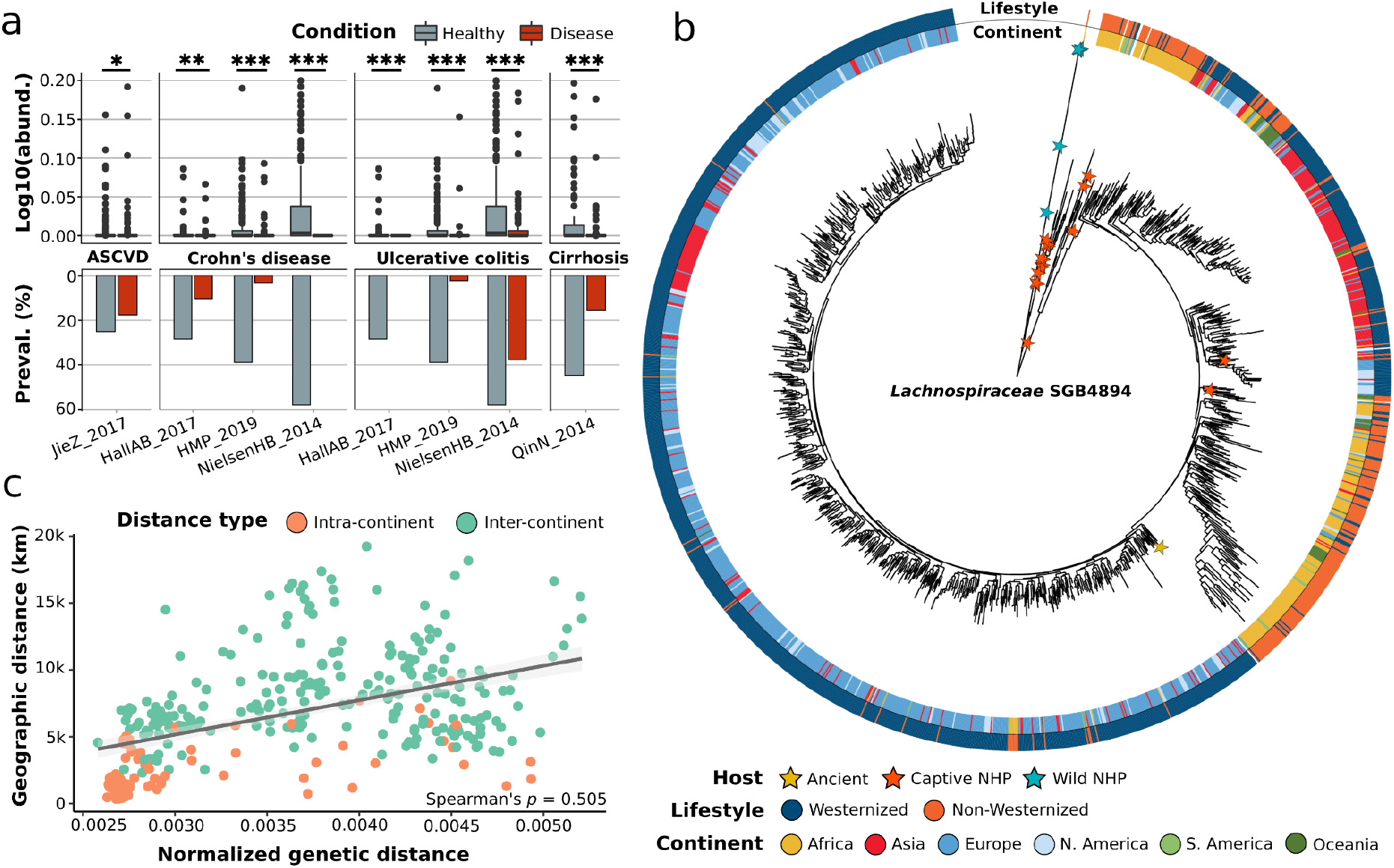
StrainPhlAn 4 accurately reconstructs large-scale strain-level phylogenies of uncharacterized microbial species. **a**, Relative abundances (box plots and top-part *y* axis) and prevalences (bar plots and bottom-part *y* axis) of the uncharacterized species (uSGB) *Lachnospiraceae* SGB4894 are significantly higher in healthy individuals in comparison with patients suffering several gastrointestinal-related diseases, and this difference is reproducible across populations (Mann-Whitney U test; *=p-value<0.05. **=p-value<0.01, ***=p-value<0.001). **b**, *Lachnospiraceae* SGB4894 shows within-species genetic diversity strongly linked to geographic origin and lifestyle. **c**, Pairwise geographic distances between strains of different countries correlate with their median genetic distances (Spearman’s *p=*0.505, see **Methods**), suggesting that human *Lachnospiraceae* SGB4894 strains could have followed an isolation-by-distance pattern.

### StrainPhlAn 4 reconstructs large phylogenies of uncharacterized microbial species

The unique clade-specific marker genes exploited by MetaPhlAn to detect and quantify microbial taxa can also be used to reconstruct the sample-specific genetic make-up of individual strains with the StrainPhlAn approach ^4,65^. MetaPhlAn 4 also extends StrainPhlAn 4 to be applicable to SGBs, and thus to uncharacterized species (uSGBs). StrainPhlAn 4 uses the MetaPhlAn 4 mapping of reads against markers to produce per-sample genotypes for the dominant strains per species (for all SGBs with sufficient coverage). Compared to StrainPhlAn 3, we improved the procedure to select and process markers and samples with a more robust and validated set of default parameters and a more stringent gap trimming strategy. We also exploit the larger marker’s database of more phylogenetically consistent SGBs (avg 189 ± 34 markers per SGB). This resulted into more accurate phylogenies compared to the previous version, with an average of 1.33% increase in correlation between StrainPhlAn phylogenetic distances and MAG-based phylogenies built on the fraction of samples in which high-quality MAGs could be reconstructed (evaluation done on 100 samples for the three most prevalent kSGBs with consistent MetaPhlAn 3 species, **Table S17** and **Supplementary Fig. 10a-f**, see **Methods**).

To illustrate the potential of StrainPhlAn profiling for uSGBs, we continued our exploration of the health-linked *Lachnospiraceae* SGB4894 introduced above, exploiting the same collection of 19.5k gut metagenomic samples used for MetaPhlAn 4 (**Table S12**). This analysis incorporated all 5.8k samples in which MetahlAn 4 detected *Lachnospiraceae* SGB4894, including 79 non-human primates and 12 ancient human gut metagenomes (**Table S18**). StrainPhlAn 4 retained 37 SGB4894-specific marker genes (spanning 19,449 nucleotide positions after trimming the alignment to exclude non-variable positions) across the 1,683 samples in which the target uSGB had enough coverage for strain profiling (samples with, at least, 20 *Lachnospiraceae* SGB4894 markers reconstructed with >80% breadth of coverage) and automatically built a phylogeny integrating all strain profiles from among host types.

The resulting phylogeny showed that *Lachnospiraceae* SGB4894 is composed of multiple sub-clades, including one comprising strains mostly from individuals from Westernized populations and other two instead dominated by individuals from non-Westernized or Chinese populations, the latter also with higher intra-clade diversity (**Fig. 5b**). One strain reconstructed from a sample of palaeofaeces from ∼1,300 years ago ^66^ was also integrated within the *Lachnospiraceae* SGB4894 phylogeny and placed as basal for the subclade of mostly European and North-American strains (**Fig. 5b**), whereas the strains from non-human primates tended to populate a common, divergent region of the tree.

*Lachnospiraceae* SGB4894’s phylogeny further demonstrated genetic structure linked to the geographic origin of the hosts (**Fig. 5b**). Indeed, when considering pairs of strains sampled in different countries, we found a remarkable correlation between geographic and median genetic distance (Spearman’s *p*=0.505), suggesting that the population genetic structure of *Lachnospiraceae* SGB4894 could have been, in part, affected by isolation-by-distance effects ^67^, as previously shown for *Helicobacter pylori* ^*68*^ and *Eubacterium rectale* ^69^ (**Fig. 5c**). Correspondingly, SGB4894 had a higher intra-population genetic variability in non-Westernized populations (Mann-Whitney U test, p-value<2.22e-46, **Supplementary Fig. 9b**) as well as higher intra-subject polymorphism rates (calculated as the percentage of bases in the reconstructed markers with an allele dominance below 80%, Mann-Whitney U test, p-value=8.6e-14, **Supplementary Fig. 9c**). StrainPhlAn 4 thus readily enabled phylogenetic reconstruction and population genetics for uncultivated, yet-to-be-named species with high precision (see **Methods, Table S17** and **Supplementary Fig. 10g**,**h**).

StrainPhlAn 4 also allows the analysis of strain sharing and transmission between communities ^4^ for uncharacterized species, i.e. uSGBs (see **Methods**). Notably, for Lachnospiraceae SGB4894, StrainPhlAn 4 estimated that strains of this uSGB were not shared between mothers and their <1 year infants in all 21 cases in which it was reliably detected in both relatives (**Supplementary Fig. 9d**). Similarly, only 5.63% of adults in the same household that were positive for *Lachnospiraceae* SGB4894 shared the same strain (**Supplementary Fig. 9d**), suggesting that stable vertical and horizontal transmission for this species are both rare. There is some evidence for horizontal transmission between host species, however, as we found evidence of two captive non-human primates sharing closely-related *Lachnospiraceae* SGB4894 strains with humans (**Fig. 5b**). Overall, this example shows that the extension of StrainPhlAn 4 to incorporate SGBs alongside MetaPhlAn 4 enables the analysis of highly-resolved, sub-species phylogenies for both well-characterized and yet-to-be-cultivated microbial species.

## Discussion

MetaPhlAn 4 provides a strategy for integrating metagenomic assembly with reference-based profiling approaches to achieve the best of both worlds: novelty by incorporating diverse high-quality metagenome assemblies, and sensitivity and specificity using refined mapping to pre-screened marker sequences. This efficiently leverages multiple recent large efforts in metagenomically cataloging microbial diversity ^37–46^, organizing over 1M prokaryotic sequences into species-level genome bins and efficiently using them to profile new metagenomes using a marker-based strategy. This quantifiably improved the resolution of health-associated biomarkers, microbial phylogeny, and uncharacterized taxa across tens of thousands of shotgun metagenomes spanning dozens of distinct environments.

Notably, even with the extended MetaPhlAn 4 SGB and marker set, further work remains to better-profile under-characterized habitats. Environmental, non-host-associated, and other under-studied microbial communities are still highly enriched for sequences not captured even by current uSGBs, although the algorithm and software architecture can be continuously updated as new MAGs become available. The current methods also do not extensively incorporate viral or eukaryotic microbial sequences, due to their unique genomic architectures and quality control requirements relative to bacterial and archaeal genomes. Interestingly, since SGBs represent essentially whole-genome operational taxonomic unit (OTU) clusters ^70^, many related downstream statistical challenges also remain to be addressed; for example, the tradeoff between sensitivity and specificity when applying quality control measures to identify real but rare taxa. We expect to continue addressing these challenges in future versions of the methodology, which will also form the basis for other MAG-aware updates of the bioBakery platform ^4,71^.

## Methods

### Overview of the approach

MetaPhlAn 4 taxonomic profiling relies on detecting the presence and estimating the coverage of a collection of species-specific marker genes to estimate the relative abundance of known and unknown microbial taxa in shotgun metagenomic samples. Since version 4, MetaPhlAn is relying on the concept of sequence-defined species-level genome bins (SGBs)

^42^ that addresses many limitations of manual taxonomy assignment and encompasses taxonomic units both with available reference genomes from cultivation (kSGBs) and taxa defined solely based on metagenome-assembled genomes (uSGBs).

As a brief summary of the approach (details in the following subsections), in order to build the MetaPhlAn database of SGB-specific markers, we collected a catalog of 729,195 dereplicated and quality-controlled genomes (560,084 MAGs and 169,111 reference genomes), that was used to expand the SGB organization by Pasolli et al. ^42^. This led to the definition of 21,373 family-level genome bins (FGBs), 47,643 genus-level genome bins (GGBs) and 70,927 SGBs, with 23,737 of them containing at least one reference genome (kSGBs) and 47,190 containing only MAGs (uSGBs). To minimize the chance that SGBs incorporate assembly artifacts or chimeric sequences, we considered only those uSGBs with at least 5 MAGs (no filtering for kSGBs). The genome catalog was then annotated using the UniRef90 database ^55^ (see below) and, within each SGB, the genes that could not be assigned to UniRef90 gene families were *de-novo* clustered together using the UniClust90 ^56^ criteria (>90% identity and >80% coverage of the cluster centroid). Using the resulting UniRef- and UniClust90 annotations, we defined a set of core genes for each quality-controlled SGB (genes present in almost all genomes composing a SGB), and after mapping all core genes against the entire genomic catalog, we defined a set of 5.1M SGB-specific marker genes (core genes not present in any other SGB) for a total of 21,978 kSGBs and 4,992 uSGBs.

For the taxonomic profiling step that uses the markers based on the SGB data, MetaPhlAn 4 maps metagenomic reads (preferably already quality-controlled) against the marker database using Bowtie2 ^72^. From these mapping results, MetaPhlAn estimates the coverage of each marker and computes the clade’s coverage as the robust average of the coverage across the markers of the same clade. Finally, the clade’s coverages are normalized across all detected clades to obtain the relative abundance of each taxa. Several downstream analyses are included in the MetaPhlAn package, including the strain-level phylogenetic profiling of SGBs by StrainPhlAn.

### The starting catalog of reference genomes and MAGs

Starting from the original catalog of 154,724 human metagenomic-assembled genomes (MAGs) and 80,990 reference genomes collected by Pasolli et al ^42^, we retrieved an additional set of 616,805 MAGs spanning different human body sites, animal hosts and non-host associated environments **(Table S2**), as well as 155,767 new reference genomes available as of November 2020 in the NCBI Genbank database ^73^. In order to ensure the quality of the downloaded sequences, we executed CheckM version 1.1.4 ^74^ on the complete catalog of 1,008,148 genomes (i.e. reference sequences and MAGs), filtering those with completeness below 50% or contamination above 5%. In order to avoid multiple inclusions of the same strains, we computed the all-versus-all MASH distances ^54^ on the quality-controlled sequences, followed by the dereplication at 99,99% genetic identity. This resulted in a quality-controlled catalog of 729,195 genomes, comprising 560,084 MAGs and 169,111 reference genomes.

### Building the expanded SGB catalog

Using the new genomic catalog, we expanded the species-level genome bin (SGB) organization proposed by Pasolli et al. ^42^. First, we apply the “*phylophlan_metagenomic”* subroutine of PhyloPhlAn 3 ^75^ on the 493,482 new MAGs and reference genomes in order to identify their closest SGB, genus-level genome bin (GGB), and family-level genome bin (FGB) and their MASH distances. Based on the reported distances, we assigned the genomes to the already existing SGBs, GGBs and FGBs according to the thresholds defined by Pasolli et al. (5%, 15%, and 30% genetic distance, respectively) ^42^. We then applied a hierarchical clustering with average linkage on the all-versus-all MASH distances of the genomes not assigned to any existing SGB, using the “fastcluster” python package version 1.1.25. The resulting dendrogram was divided with cutoffs at 5%, 15% and 30% genetic distance to define 54,596 new SGBs, 37,546 new GGBs and 18,211 new FGBs. In short, from the initial filtered catalog of 729,195 MAGs and reference genomes, we defined 21,373 FGBs, 47,643 GGBs and 70,927 SGBs, with 23,737 of them containing, at least, one reference genome (kSGBs) and 47,190 containing only MAGs (uSGBs) (**Table S1**).

We assigned a taxonomic label to all the 70,927 SGBs according to the NCBI taxonomy database (as of February 2021) ^49^. For kSGBs, we assigned the taxonomy by applying a majority rule to the taxonomic labels of the reference genomes contained in each SGB. In case of a tie, the taxonomic label will be resolved by choosing as the representative taxon the one alphabetically first. For uSGBs, we applied a similar majority rule but on the taxonomies of the reference genomes contained at the GGB level, assigning the taxonomic label up to the genus level. If no reference genomes were present at the GGB level, we further applied the same procedure at the FGB level. If no reference genomes were found at the FGB level, we assigned the taxonomic labels only up to the phylum level by considering the phylum that is most recurrent within the set of taxonomic labels of the closest reference genome and, at most, up to one hundred reference genomes within 5% genomic distance to the closest as identified by “*phylophlan_metagenomic*”. For the taxonomic levels not receiving any taxonomy label, we assigned all the internal taxonomic nodes with SGB, GGB, and FGB identifiers to maintain the taxonomy with all its levels and for providing categorization of uSGBs.

### Genome annotation and pangenome generation

The filtered catalog of 729,195 MAGs and reference genomes was subjected to an annotation workflow in which (i) the FASTA files were processed with Prokka (version 1.14) ^76^ in order to detect and annotate the coding sequences (CDS) and (ii) subsequently assign the CDS to a UniRef90 cluster ^55^ using a DIAMOND-based pipeline (available in https://github.com/biobakery/uniref_annotator). The DIAMOND-based pipeline, performs a sequence search (DIAMOND version 0.9.24) ^77^ of the protein sequences against the UniRef90 database (release 2019_06) and then applies the UniRef90 inclusion criteria on the mapping results to annotate the input sequences (>90% identity and >80% coverage of the cluster centroid). Within each SGB, protein sequences that were not assigned to any UniRef90 cluster were clustered using MMseqs2 ^78^ following the Uniclust90 criteria (“*-c 0.80 --min-seq-id 0.9”* parameters) ^56^.

For each SGB, based on the UniRef90 and UniClust90 annotations, a pangenome was generated by collecting all the UniRef/UniClust90 clusters present in at least one of the SGB’s genomes. For each cluster, the representative sequence was randomly selected within all the genomes and a coreness value was calculated based on the cluster prevalence within the 2k highest quality genomes of the SGB. uSGBs containing less than 5 MAGs were discarded for the following steps. We implemented this restriction since we found evidence that some of the small uSGBs contained likely assembly artifacts or chimeric genomes. In this step, 41,498 uSGBs out of the 70,927 SGBs were discarded, while all kSGBs were retained as they are represented by theoretically more reliable sequences.

### The MetaPhlAn 4 vJan21 markers database

From these pangenomes, the construction of the marker database for MetaPhlAn 4 is divided in two sequential steps: the identification of the core genes within each SGB and the screening of the core genes for their SGB-specificity.

For the identification of the core genes, the procedure first defines a coreness percentage threshold (i.e. the percentual prevalence of a gene within the SGB) based on the SGB pangenomes. Specifically, we selected the maximum coreness threshold that allowed the retrieval of at least 800 core genes (of length between 450 and 4,500 nucleotides). The minimum coreness threshold was bound to 60% for SGBs with less than 100 genomes, and to 50% for the others. For each SGB, a core genes set was generated using the inferred coreness thresholds. On average, we retrieved 2,985 core genes per SGB (median 2,687, s.d. 1,861). SGBs with less than 200 core genes were discarded and not considered further (9 SGBs).

In order to detect the SGB-specific marker genes, each set of core genes was then aligned against the genomes of the other SGBs using Bowtie 2 (version 2.3.5.1; *--sensitive* parameter) ^72^. For each SGB, a subset comprising up to the highest quality 100 genomes was selected for the mapping for computational reasons. Each core gene was split in fragments of 150 nt length in order to simulate metagenomic reads, and then they were mapped against the representative subset of the SGB’s genomes. An alignment hit of a fragment was considered a hit for its corresponding core gene. Core genes hitting none (perfectly unique markers) or less than 1% (quasi-markers) of the genomes of any other SGB and hitting a number of the genomes of their SGB above or equal to their coreness threshold were selected as marker genes. Crucially, this uniqueness procedure was substantially stricter than those used in previous MetaPhlAn versions owing to the improved consistency of the SGBs compared to original species taxonomic assignments.

The small fraction of SGBs producing less than 100 marker genes (810 SGBs) were subjected to the following workflow:

1. If more than 200 core genes of the target SGB were matching an external SGB (a kSGB belonging to the same species, or a uSGB) and if the external SGB had less than 10% of the genomes in the target SGB, then the external SGB was discarded (this occurred for 392 kSGBs and 150 uSGBs). This step was repeated every time an external SGB was removed until the target SGB produced 100 marker genes or there were no more external SGBs that could be evaluated. In the latter case, the removal of the external SGBs was rolled back.
2. If the target SGB still could not identify 10 marker genes, external SGBs with low-quality species taxonomic labels were discarded (this occurred for 822 kSGBs and 286 uSGBs). Specifically, the regular expressions used to detecting low-quality species taxonomic labels are:
*“ (C*|*c)andidat(e*|*us)* | *_sp(_.\**|*$)* | *(.*_*|*^)(b*|*B)acterium(_.\**|) | .*\*(eury*|*)archaeo(n_*|*te*|*n$).\** | .*\*(endo*|*)symbiont.\** | .*\*genomosp_.\** | .*\*unidentified.\** | .*\*_bacteria_.\** | .*\*_taxon_.\** | .*\*_et_al_.\** | .*\*_and_.\** | .*\*(cyano*|*proteo*|*actino)bacterium_.*)*
This step was repeated every time an external SGB was removed until the target SGB produced 10 marker genes or there were no more external SGBs that could be evaluated. In the latter case, the removal of the external SGBs was rolled back.
3. For the SGBs that still did not produce at least 10 marker genes, a conflict graph was generated collecting all the core gene hits against external SGBs in which more than 200 core genes were in conflict. The graph was then processed by merging SGBs with a procedure that minimizes the number of merged SGBs and maximizes the number of markers retrieved. After this process, 849 SGBs were merged, producing 237 SGB groups.

Finally, for each SGB, a maximum or 200 marker genes were selected based first on their uniqueness and then on their size (longer first). SGBs that still had fewer than 10 markers were discarded (188 SGBs). Each marker was associated with an entry in the MetaPhlAn 4 vJan21 database which includes the SGB for which the sequence is a marker, the list of SGBs sharing the marker, the sequence length, and the taxonomy of the SGB. This produced a list of 5.1M marker genes for a total of 21,978 kSGBs and 4,992 uSGBs.

### MetaPhlAn 4 taxonomic profiling

MetaPhlAn 4 taxonomic profiling is based on read homology to and coverage of SGB-specific markers to estimate the relative abundance of taxonomic clades present in a metagenomic sample. The MetaPhlAn pipeline starts by mapping the raw reads of metagenomic samples against the SGB-specific markers’ database using Bowtie 2 ^72^. Input metagenomic reads can be provided as a single FASTQ file (compressed with several algorithms), multiple FASTQ files included in a single (compressed) archive, or as a pre-performed mapping (*bowtie2out* format). By default, the Bowtie2 mapping is performed using the “--*very-sensitive”* preset. For read-mapping quality purposes, short reads (reads shorter than 70bp; “*--read_min_len*” parameter) and low-quality alignments (alignments with a MAPQ value lower than 5; “*--min_mapq_val*” parameter) are discarded.

Using the quality-controlled mapping results, MetaPhlAn estimates the coverage of each marker and computes the clade’s coverage as the robust average of the coverage across the markers of the same clade, but excluding the top and bottom quantiles of the marker abundances (*“--stat_q”* parameter). For the SGB profiling, this parameter is by default set to 0.2, thus excluding the 20% of markers with the highest abundance as well as the 20% of markers with the lowest abundance. The coverage of quasi-markers is not considered from this computation when at least 33% (default value, *“--perc_nonzero*” parameter) of the markers of their respective external SGB were present. The clade’s coverages are finally normalized across all detected clades to obtain the relative abundance of each taxa as previously described ^2,3^.

### MetaPhlAn 4 unclassified reads calculation

MetaPhlAn 4 includes a feature for estimating the fraction of input reads that cannot be assigned to taxa in the database (*“--unclassified_estimation*” parameter). This is calculated by subtracting from the total number of input reads the average read depth of each reported SGB normalized by its SGB-specific average genome length as follows:

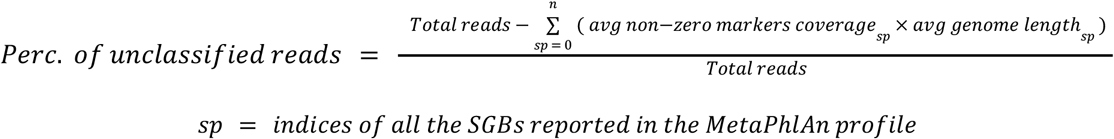

The average read depth of a SGB is calculated as the mean read depth of all its detected (non-zero) marker genes. The SGB-specific genome length for kSGBs is calculated using only the genome lengths of its reference genomes, while for uSGBs the average genome length is incremented by 7% (calculated to be the average difference between the genome sizes of references genomes and MAGs within the same SGB).

### MetaPhlAn 4 synthetic evaluation

We evaluated MetaPhlAn 4 using different published and newly created synthetic metagenomes. Firstly, we assessed the performance of MetaPhlAn 4 in comparison to several available alternatives, i.e. MetaPhlAn 3 ^4^, mOTUs 2.6 ^6^ and Bracken 2.5 ^5^. Through the OPAL benchmarking framework ^58^, we evaluated the performance of each tool by profiling the CAMI II taxonomic profiling challenge metagenomes ^57^ and SynPhlAn-nonhuman synthetic metagenomes ^4^. The CAMI II metagenomes include 118 samples representing five human body site-specific microbiomes (i.e. airways, oral, the gastrointestinal tract, skin and the urogenital tract) and the murine gut microbiome, while the SynPhlAn-nonhuman metagenomes were designed to mirror the sequencing depth and community structure of the CAMI II metagenomes (i.e. 30 million, 150-nt paired-end sequencing reads from genomes in kSGBs with a log-normal abundance distribution), but for environments different than the human body.

We ran each tool using default parameters with the exception of mOTUs 2.6, which was run twice with parameters “*-C recall*” and “*-C precision*” in order to optimize for precision and recall separately, respectively. Additionally, in order to better evaluate the SGB architecture, we performed an alternative evaluation assessing the detection and quantification of the genomes included in the synthetic metagenomes. To this end, we defined (i) “true positive” as the detection of an SGB containing a genome present in the synthetic metagenome, (ii) “false positive” as the detection of an SGB that do not contain any genome in the metagenome, and (iii) “false negative” as the non-detection of a SGB containing a genome present in the synthetic metagenome. Detection of an SGB that represents an overlapping SGB present in the community was also accounted as “true positive”. For the gold standard, relative abundances were obtained by summing-up the relative abundances of the genomes belonging to the same SGB. For MetaPhlAn 3 that contains markers describing species groups, we considered as (i) “true positive” a species group containing a species present in the synthetic metagenome, and as (ii) “false positive” a species group that do not contain any species present in the synthetic metagenome.

In order to further assess the performance of MetaPhlAn 4 to profile both known and unknown SGBs complementing the synthetic samples from CAMI II and SynPhlAn, we constructed additional synthetic metagenomes from different environments, hosts and human body sites using ART ^79^ with the Illumina HiSeq 2500 error model (available at http://segatalab.cibio.unitn.it/tools/metaphlan/). For each environment, we simulated 5 metagenomes containing 30 million, 150-nt paired-end sequencing reads using randomly-selected genomes from SGBs containing MAGs coming from that environment (with a restriction of one genome per SGB), and following a log-normal abundance distribution. MetaPhlAn 4 evaluation was then performed by assessing the detection and quantification of the genomes included in the synthetic metagenomes as described above. Additionally, to demonstrate that the evaluation was not biased by the usage of genomes included in the genomic catalog, we built, using the same procedure, another 5 metagenomes using a mixture of novel MAGs and reference genomes not included in our genomic database. SGB assignment of the new genomes was performed using the “*phylophlan_metagenomic”* subroutine of PhyloPhlAn 3 ^75^ against the Jan21 database.

### MetaPhlAn 4 application to human and non-human metagenomes

To measure the increase of the fraction of classified reads when compared with MetaPhlAn 3, we profiled 24,515 samples from 145 datasets spanning different human body sites (airways, gastrointestinal tract, oral, skin and urogenital tract) and lifestyles, animal hosts (non-human primates, mice and ruminants) and other non-host associated environments (soil, freshwater and ocean) (**Table S8**) with both MetaPhlAn 3 (version 3.0.12) and MetaPhlAn 4 (version 4.beta.1) using the unknown/unclassified estimation feature (**Table S19**). Improvements were reported using only the samples in which both tools reported, at least, one species. A SGB was reported to be present in a specific environment if it was detected in, at least, 1% of the samples from that environment. Finally, to investigate the abundance and prevalence of gut-related SGBs across different age categories and lifestyles, we selected a subset of 19,468 human gut metagenomes from 86 datasets for which age information was available (**Table S12**) as reported and curated in the curatedMetagenomicData ^80^ package.

### Westernization definition

The process of Westernization brought by industrialization and urbanization over the past two hundred years has had a significant impact on human populations. These changes include access to pharmaceuticals and healthcare, improved sanitation and hygiene, increased urban dwelling and decreased exposure to livestock, as well as changes in habitual diets (with Westernized diets tending to consist of increased fat and animal proteins, high salt and simple carbohydrates). In this study we characterize Westernized or non-Westernised individuals or populations based on either the distinction given in the primary publication or an assessment based on the criteria outlined above.

### Differential abundance analysis of diet-related taxa in the mouse microbiome

We performed a differential abundance analysis of high-fat (HF) versus normal chow diets in a public cohort of 181 murine gut microbiome ^61^. From the original cohort, we excluded 10 samples missing age information, and we selected only the samples coming from genetic backgrounds tested for both types of diet. In total we analyzed 43 HF-fed mice and 88 mice fed with normal control chow, further stratified into 2 genetic backgrounds and 5 vendors (**Table S14**). To correct for data compositionality, we first imputed the zero-values with the minimum value for an abundance found in the dataset, then we applied the centered-log-ratio transformation to the SGB’s relative abundance distribution (“*scikit-bio*” Python package, version 0.5.6). For each feature (SGB), we then built a random-intercept model using the “*statsmodels*” Python package version 0.11.1. We associated the diet (HF or chow, encoded as a binary factor) to the transformed abundance of the strain, using the sex, age-in-days and genetic background of the mice as fixed effects, and the vendor as grouping variable. Significance was assessed via Wald-test. P-values were Benjamini-Hochberg corrected (“*statsmodels*” Python package, Q<0.2). Before plotting, we selected the biomarkers having a mean abundance in the associated group greater than 1%. The reported heatmap was printed using the “*pheatmap*” R package version 1.0.12 (parameters “*clustering_distance_cols = ‘euclidean’, clustering_method = ‘complete’, cluster_rows = FALSE*”).

### Associations between microbiome and host diet and cardiometabolic markers

We assessed the associations between microbiome and cardiometabolic health and dietary patterns using 1,001 deeply phenotyped individuals from the UK retrieved from the ZOE PREDICT 1 intervention study ^22^. Machine learning (ML) analyses were performed using the “*scikit-learn*” Python package (version 0.22.2) on a panel of 19 representative nutritional and cardiometabolic markers described in the original study ^22^. A cross-validation approach was implemented using an 80/20 random splitting of training and testing sets, repeated for 100 bootstrap iterations, again with the same exact approach of the original study. Since the ZOE PREDICT 1 cohort includes twins, to avoid overfitting the twin from the training set was removed if its twin pair was present in the testing set. The ML model is based on random forests (RFs) using SGBs-level taxonomic relative abundances as estimated by MetaPhlAn 4 and relative abundance values were arcsin-sqrt transformed.

For the RF classification task, continuous features were separated into two classes, the top and bottom quartiles. The “*RandomForestClassifier”* function was used with parameters “*n_estimators=1000, max_features=‘sqrt’*”. For the RF regression task, the RandomForestRegressor function was used with parameters “*n_estimators=1000, criterion=‘mse’, max_features=‘sqrt’*”. A linear regressor (“*LinearRegression”* function with default parameters) was also trained on training target values to calibrate the range of output values predicted by the RF regressor model. Pairwise Spearman’s correlations were computed between the relative abundance of uSGBs with a prevalece of at least 20% (at least 200 of 1001 samples), and the panel of 19 nutritional and cardiometabolic markers, correcting for age, sex and body mass index. Correlations were computed using the “*ppcor*” R package version 1.1 (**Table S20**) and p-values were corrected through the Benjamini-Hochberg procedure.

### *Lachnospiraceae* SGB4894 association with host health conditions

To investigate associations between *Lachnospiraceae* SGB4894 with host health conditions across several diseases, we collected 21 disease case/control datasets available through curatedMetagenomicData ^80^ (**Table S16**). For each dataset, we assessed the associations of *Lachnospiraceae* SGB4894 with the subjects reported as healthy controls by computing a Mann-Whitney U test on the arcsin square root transformed relative abundances profiles using the “*stats.mannwhitneyu*” function of the “*scipy*” Python package version 1.5.2. Samples from Westernized adults were used and comparisons were performed only when at least 10 healthy and 10 disease samples were available. Statistically significant associations were defined by a p-value<0.05.

### StrainPhlAn 4 profiling

StrainPhlAn profiling estimates strain-level species-specific phylogenies, and it is based on the reconstruction of sample-specific consensus sequences of MetaPhlAn species-specific marker genes followed by multiple-sequence alignment and phylogenetic inference ^4,65^. Compared to StrainPhlAn 3, the accuracy and performance of StrainPhlAn 4 have been improved mostly because of (i) the redesigned procedure to select and process markers and samples to be considered in the phylogeny, and (ii) the use of the same MetaPhlAn 4 database of markers from the extensive set of phylogenetically consistent SGBs.

For item (i), StrainPhlAn 4 considers as input the reads-to-markers alignment results (in SAM format ^81^) from the MetaPhlAn 4 profiling together with the MetaPhlAn 4 database. For each sample, StrainPhlAn 4 reconstructs consensus sequences of the species-specific marker genes by considering, for each position, the nucleotide with the highest frequency among the reads mapping against it. By default, consensus markers covered by less than eight reads or with a breadth of coverage below 80% are discarded (i.e. the proportion of the marker covered by reads, “*--breadth_threshold*” parameter). For this step, ambiguous bases (i.e. positions in the alignment with quality lower than 30 or with major allele dominance below 80%) are considered as unmapped positions. After the reconstruction of the markers, StrainPhlAn discards samples with less than 80% of the available markers and markers present in less than 80% of the samples (“*--sample_with_n_markers*” and “*--marker_in_n_samples*” parameters, respectively). Then, markers are trimmed by removing the leading and trailing 50 bases (“*--trim_sequences*” parameter), and a polymorphic rates report is generated. Finally, the remaining samples and markers are processed by PhyloPhlAn ^75^. By default, multiple sequence alignment is performed by MAFFT ^82^, gappy positions (i.e. positions with more than 67% of gaps) are trimmed by trimAl ^83^, and phylogenetic trees are inferred by RAxML ^84^.

### *Lachnospiraceae* SGB4894 strain-level analyses

For the *Lachnospiraceae* SGB4894 strain-level analysis, we selected 5,883 human gut metagenomic samples from 86 datasets in which *Lachnospiraceae* SGB4894 was reported to be present based by MetaPhlAn 4 (**Table S12**). 79 non-human primates (NHP) and 12 ancient human gut metagenomic samples were also included from 12 different datasets (**Table S18**). SGB4894-specific marker genes were successfully reconstructed from 2,787 metagenomes, of which 2,738 were from contemporary human gut microbiome samples, 5 from ancient gut microbiome samples, and 44 from NHP gut microbiome samples. Strain-level profiling with StrainPhlAn 4 was performed using parameters “*--marker_in_n_samples 70 —sample_with_n_markers 10 ––phylophlan_mode accurate*”. The phylogenetic tree generated by StrainPhlAn was plotted with GraPhlAn version 1.1.4 ^85^. Phylogenetic distances were extracted based on the distance between samples in the tree and normalized by the total branch length of the tree. Geographic distances between countries were calculated using the “*distGeo*’’ function of the “*geosphere*” R package version 1.5-10. Spearman’s correlation between genetic and geographic distance was then calculated using the “*cor.test*” function of the “*stats”* R package. Finally, in order to assess the transmissibility of *Lachnospiraceae* SGB4894, we executed the StrainPhlAn’s “*strain_transmission.py*” script using as input the phylogenetic tree (default parameters). The script, which is part of the StrainPhlAn release, can use the species specific cut-offs on the normalized phylogenetic distances pre-computed on the available datasets with longitudinal sampling.

### StrainPhlAn 4 evaluation

The three most prevalent single-species kSGBs whose species were available in the MetaPhlAn 3 database, i.e. *Blautia wexlerae* (SGB4837), *Bacteriodes uniformis* (SGB1836), and *Eubacterium rectale* (SGB4933), were selected to evaluate the improvements included in StrainPhlAn 4 in comparison with the previous version. As a gold standard, for each species, we considered 100 high-quality MAGs randomly selected from the genomic catalog (**Table S21**), and obtained a phylogeny by processing the MAGs via Roary core gene alignment and RAxML tree reconstruction. Specifically, we computed a multiple sequence alignment from each set of core genes (present in at least 90% of genomes) using Roary version 3.13.0 ^86^ with parameters “*-cd 90 -i 90 -e --mafft*”, and launched RAxML version 8.2.4 ^84^ with parameters “*-f a -# 100 -p 12345 -x 12345 -m GTRGAMMA*”. Using the metagenomic samples from which the considered MAGs were assembled from, we executed StrainPhlAn 3 and 4 using their respective database and with default parameters and “*–mutation_rates*”. Additionally, we executed a similar evaluation (but using the MetaPhlAn 4 database in the StrainPhlAn 3 call) on the uSGB *Lachnospiraceae* SGB4894, using the 170 MAGs from the genomic catalog with publicly available metagenomic samples. Pairwise phylogenetic distances normalized by the total branch length were calculated using the PyPhlAn package (https://github.com/SegataLab/pyphlan). Pearson correlations between StrainPhlAn and the gold standard results were calculated using the “*stats.pearsonr*” function of the “*scipy*” Python package version 1.5.2.

## Data availability

All metagenomic studies analyzed in this work are publicly available through the corresponding publications listed in **Table S8**. The CAMI II Challenge synthetic metagenomes and gold standards are available at https://data.cami-challenge.org/participate. The SynPhlAn-nonhuman synthetic metagenomes and gold standards are available at http://segatalab.cibio.unitn.it/tools/biobakery. The novel synthetic metagenomes containing kSGBs and uSGBs and gold standards are available at http://segatalab.cibio.unitn.it/tools/metaphlan/. Prevalences of the SGBs across environments, age categories and lifestyles are available in **Tables S10-11**. Metadata of the publicly analyzed human metagenomes is also available through the curatedMetagenomicData R package ^80^. The full list of metagenomic studies used for the strain-level analysis of *Lachnospiraceae* SGB4894 is reported in **Tables S12 and S18**.

## Code availability

The MetaPhlAn 4 version described in this work is labeled as MetaPhlAn 4.beta.1 and is available at http://segatalab.cibio.unitn.it/tools/metaphlan and https://huttenhower.sph.harvard.edu/metaphlan/ with the open source code at https://github.com/biobakery/MetaPhlAn together with StrainPhlAn 4. It is also available via Bioconda (https://anaconda.org/bioconda/metaphlan) and PIP (https://pypi.org/project/MetaPhlAn).

## Acknowledgements

We would like to thank all the members of the Segata and Huttenhower lab for their insightful contributions to the work, and the users of past versions of MetaPhlAn for their suggestions and support. The work was supported by the European Research Council (ERC-STG project MetaPG-716575) to NS; by MIUR ‘Futuro in Ricerca’ (grant No. RBFR13EWWI_001) to NS; by the European H2020 program (ONCOBIOME-825410 project and MASTER-818368 project) to NS; by the National Cancer Institute of the National Institutes of Health (1U01CA230551) to NS; by the Premio Internazionale Lombardia e Ricerca 2019 to NS; by the Harvard Chan Microbiome Analysis Core (CH); by the National Institute of Diabetes and Digestive and Kidney Diseases of the National Institutes of Health (R24DK110499) to CH; by Cancer Research UK Grand Challenge award C10674/A27140 to Wendy Garrett (CH); and by the National Institute of Allergy and Infectious Diseases (U19AI110820) to David Rasko (CH).

## Contributions

ABM and NS conceived the study. ABM wrote, validated, and tested the code and performed most of the analyses. FB, FC, LJM, KNT, MZ, PM, LD, KDH, AMT, GP, EPiperni, MP, MVC, AT, EAF, EPasolli, and FA supported the development and validation of the method and of the software. FG, RD, JW, SEB, TDS contributed to the analyses and their interpretation. ABM, FA, CH, and NS wrote the paper with contribution and editing from all the authors. CH and NS supervised the work. All the authors read and approved the final version of the manuscript.

