## Supplementary Figures for "Extending and improving metagenomic taxonomic profiling with uncharacterized species with MetaPhlAn 4"

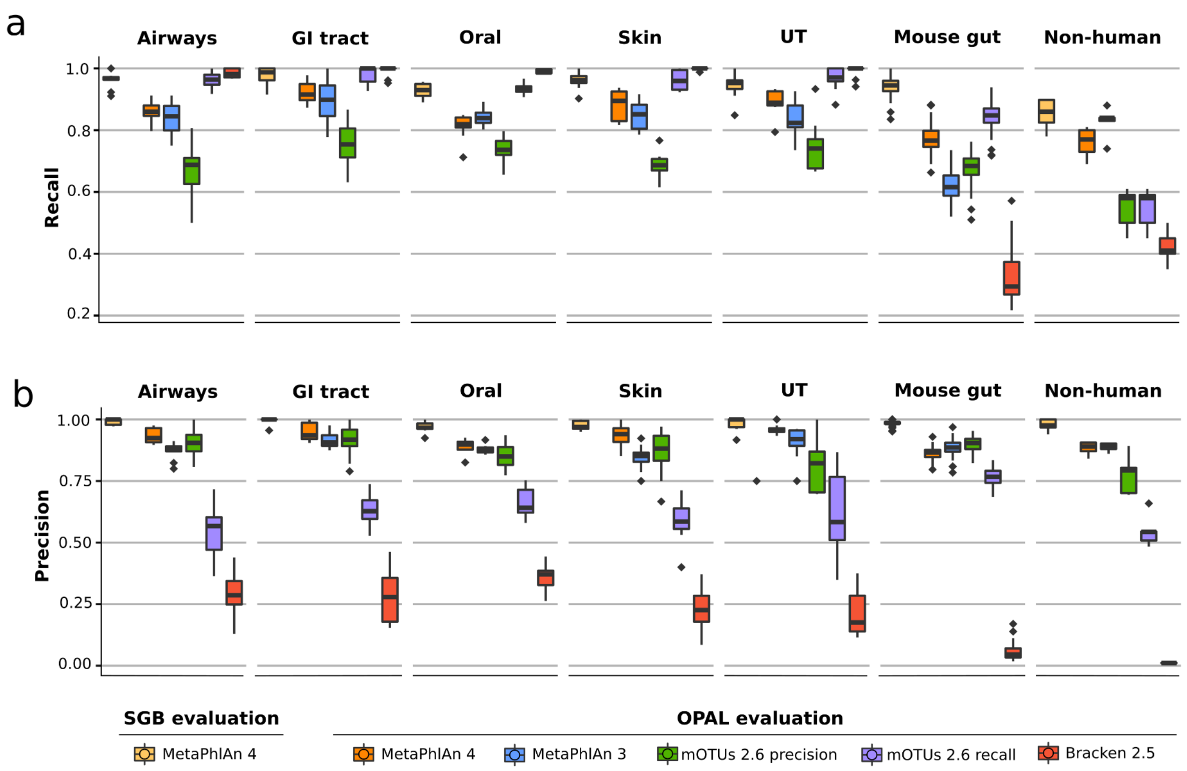


**Supplementary Figure 1. Performance of MetaPhlAn 4 using the CAMI II taxonomic profiling challenge and SynPhlAn-nonhuman synthetic metagenomes in comparison with several available alternatives.** MetaPhlAn 4 shows high accuracy when assessing (**a**) recall and (**b**) precision. GI=gastrointestinal, UT=urogenital tract.

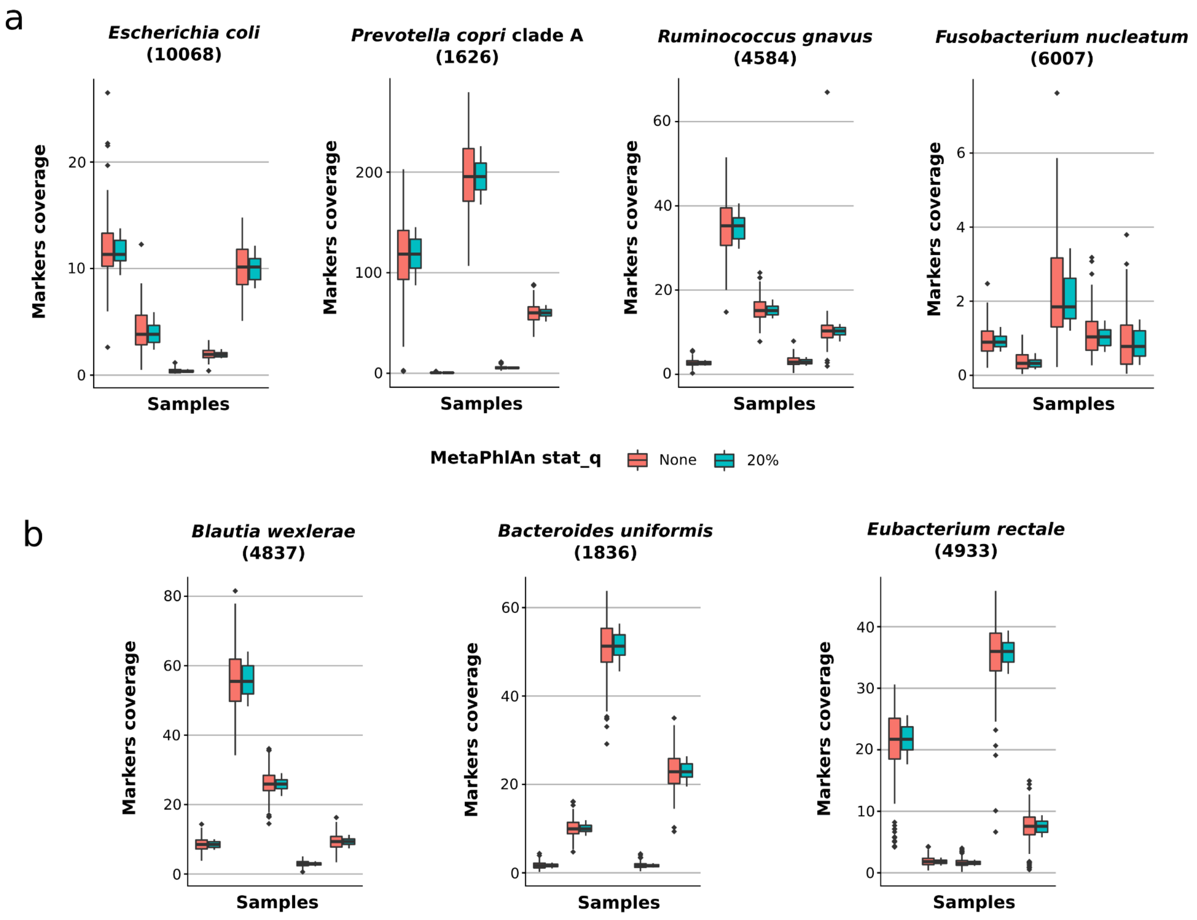


**Supplementary Figure 2. Per sample coverage of the MetaPhlAn 4 markers.** MetaPhlAn 4 markers show a high coverage consistency when assessing (**a**) biologically interesting species as well as (**b**) the three most prevalent kSGBs. Five randomly selected samples were chosen for each species evaluation. Top y-axis (red boxplots) represents the markers' coverage without applying any stat_q filtering and the bottom y-axis (blue boxplots) represents the markers' coverage while applying MetaPhlAn 4 default stat_q filtering (20%, see **Methods**).

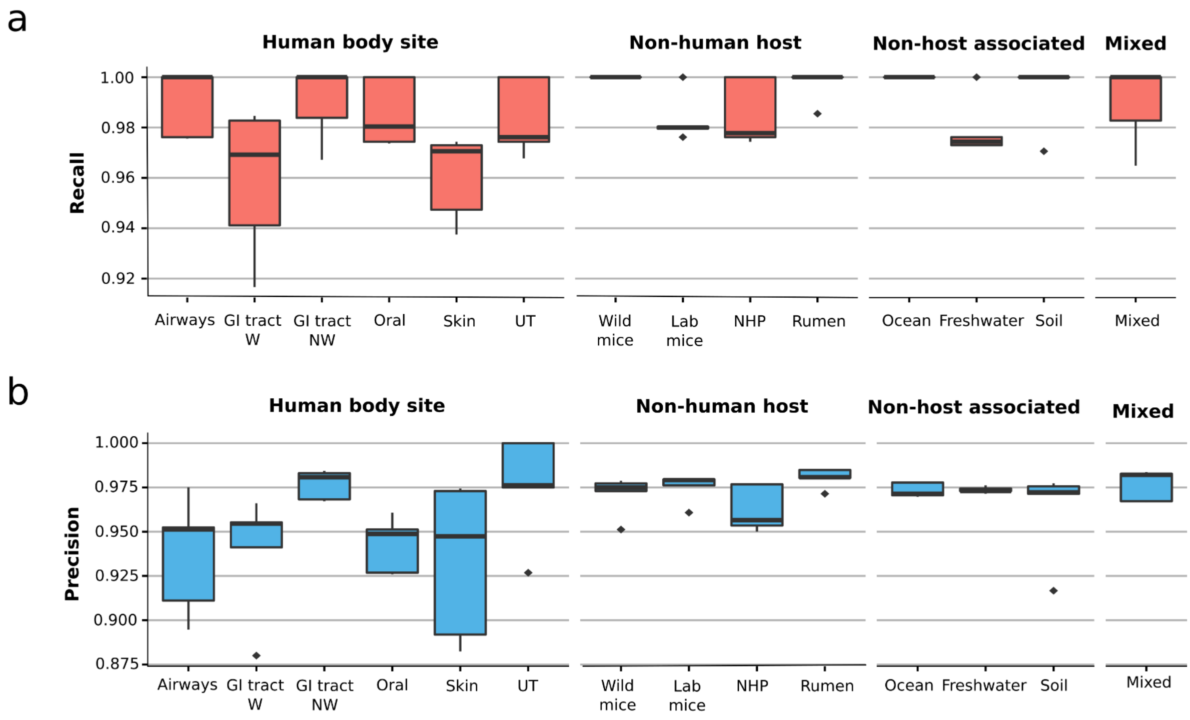


**Supplementary Figure 3. Performance of MetaPhlAn 4 using a new synthetic dataset containing both known and unknown SGBs.** MetaPhlAn 4 shows high accuracy when assessing (**a**) recall and (**b**) precision. GI=gastrointestinal, W=Westernized, NW=non-Westernized, NHP=non-human primates.

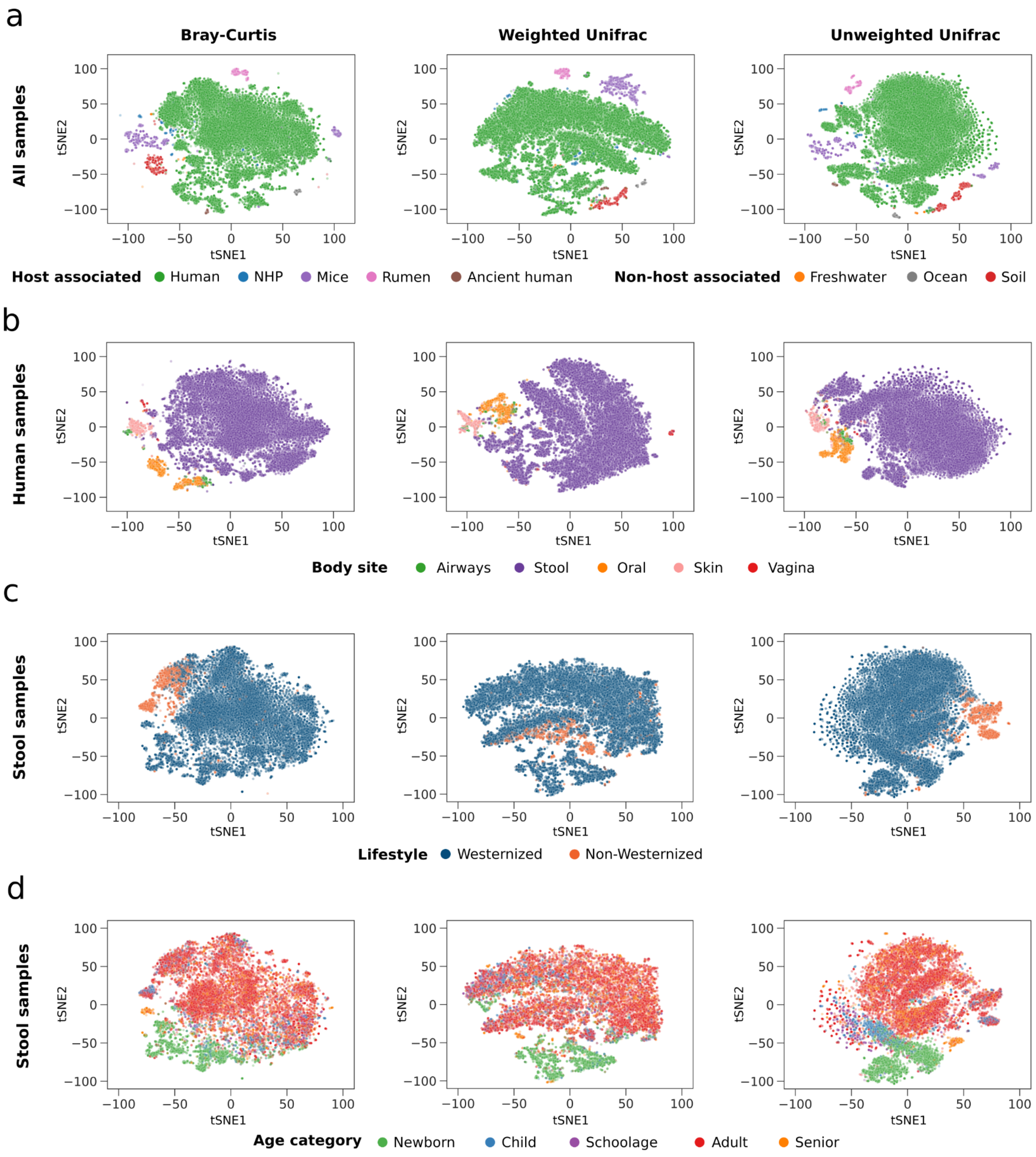


**Supplementary Figure 4. t-SNE representation of the 24.5k metagenomic samples based on (from left to right) Bray-Curtis, Weighted Unifrac and Unweighted Unifrac distances of the MetaPhlAn 4 SGB-level taxonomic profiles.** (**a**) Dimensionality reduction using all 24.5k metagenomic samples classified by environment. (**b**) Dimensionality reduction using only the 19.5k human metagenomic samples classified by body site. (**c**) Dimensionality reduction using only modern human stool metagenomic samples classified by lifestyle. (**d**) Dimensionality reduction using only modern human stool metagenomic samples classified by age category. NHP = Non-human primate.

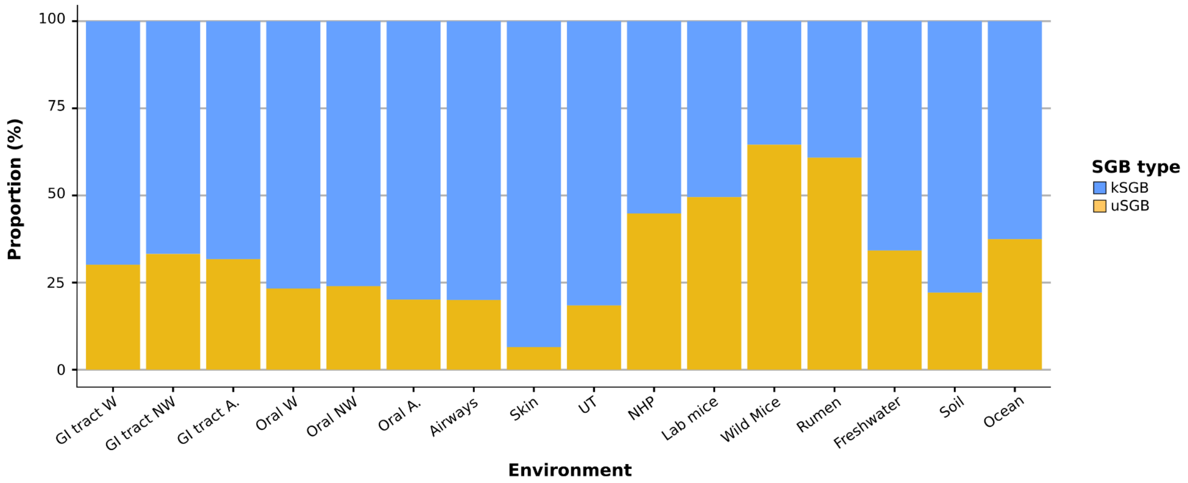


**Supplementary Figure 5. Proportion of the known and unknown SGBs across different human body sites and lifestyles, animal host and non-host associated environments**. GI=gastrointestinal, W=Westernized, NW=non-Westernized, NHP=non-human primates.

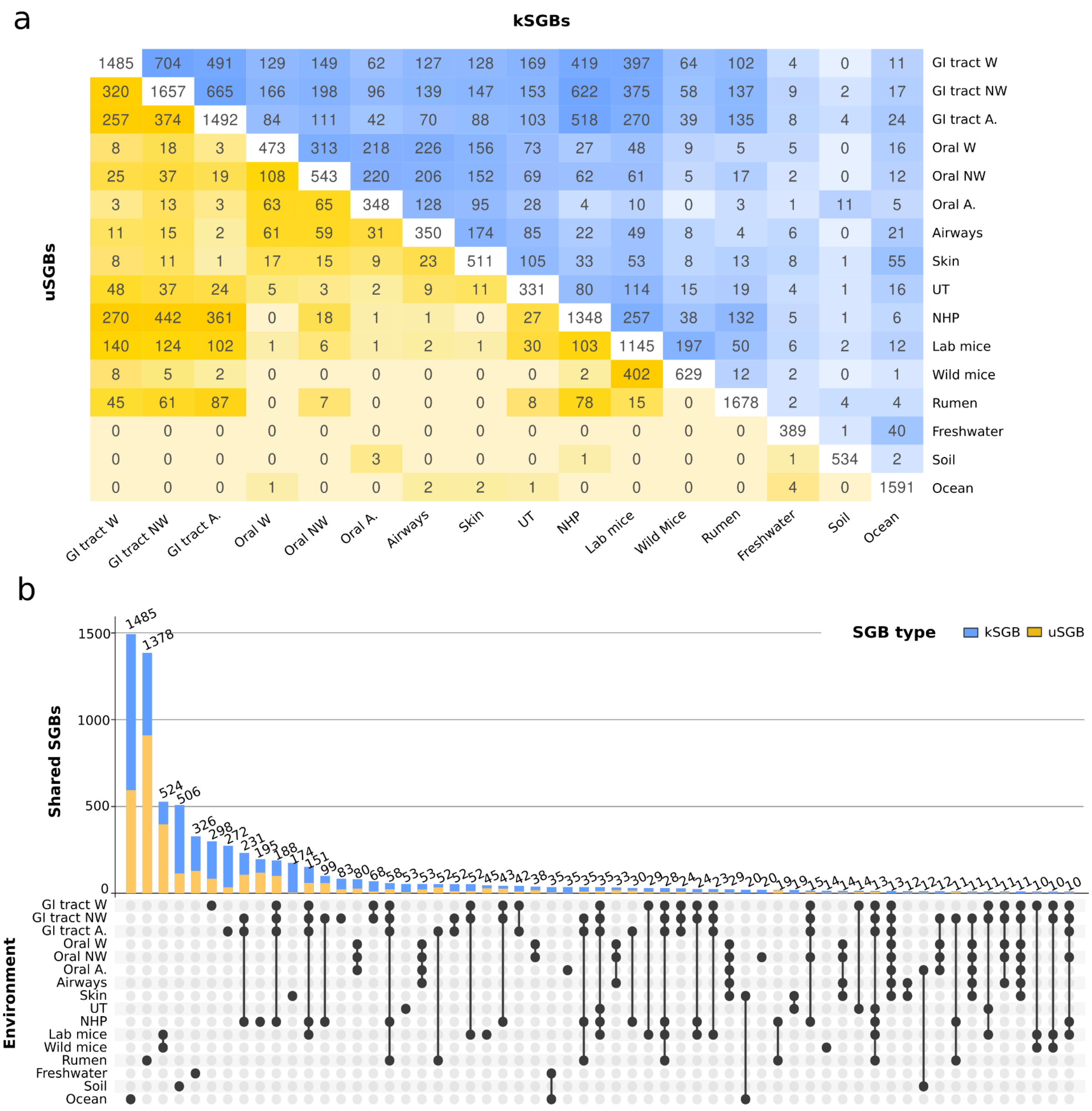


**Supplementary Figure 6. Co-presence of SGBs across several human, animal and non-host related environments**. (**a**) While a good number of microbial species are shared between the gut-associated environments, only a few and mainly characterized species are also found shared in other host- non non-host- associated environments. The top blue triangular represents the number of kSGBs shared between environments, the bottom yellow triangular shows the number of uSGBs shared between each pair of environments and the white diagonal contains the total number of SGBs detected in each individual environment. (**b**) Large portions of the SGBs only found in non-human microbiomes remain uncharacterized (uSGBs). The upset plot represents the SGBs exclusively present in each group of environments. Only intersections with more than 10 SGBs are shown. GI=gastrointestinal, W=Westernized, NW=non-Westernized, NHP=non-human primates.

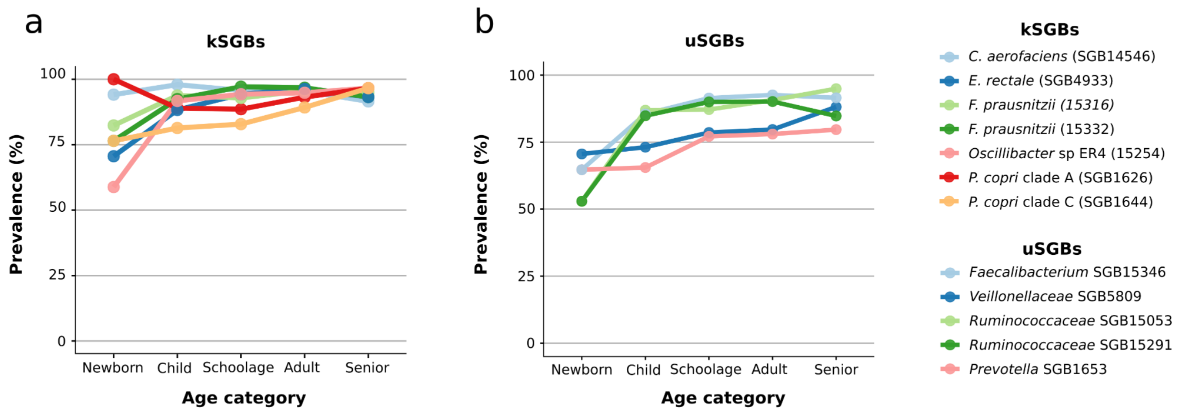


**Supplementary Figure 7. Most prevalent known (left) and unknown (right) SGBs in the Non-Westernized populations and their abundance through life.** The 2 most prevalent SGBs of each age category are shown.

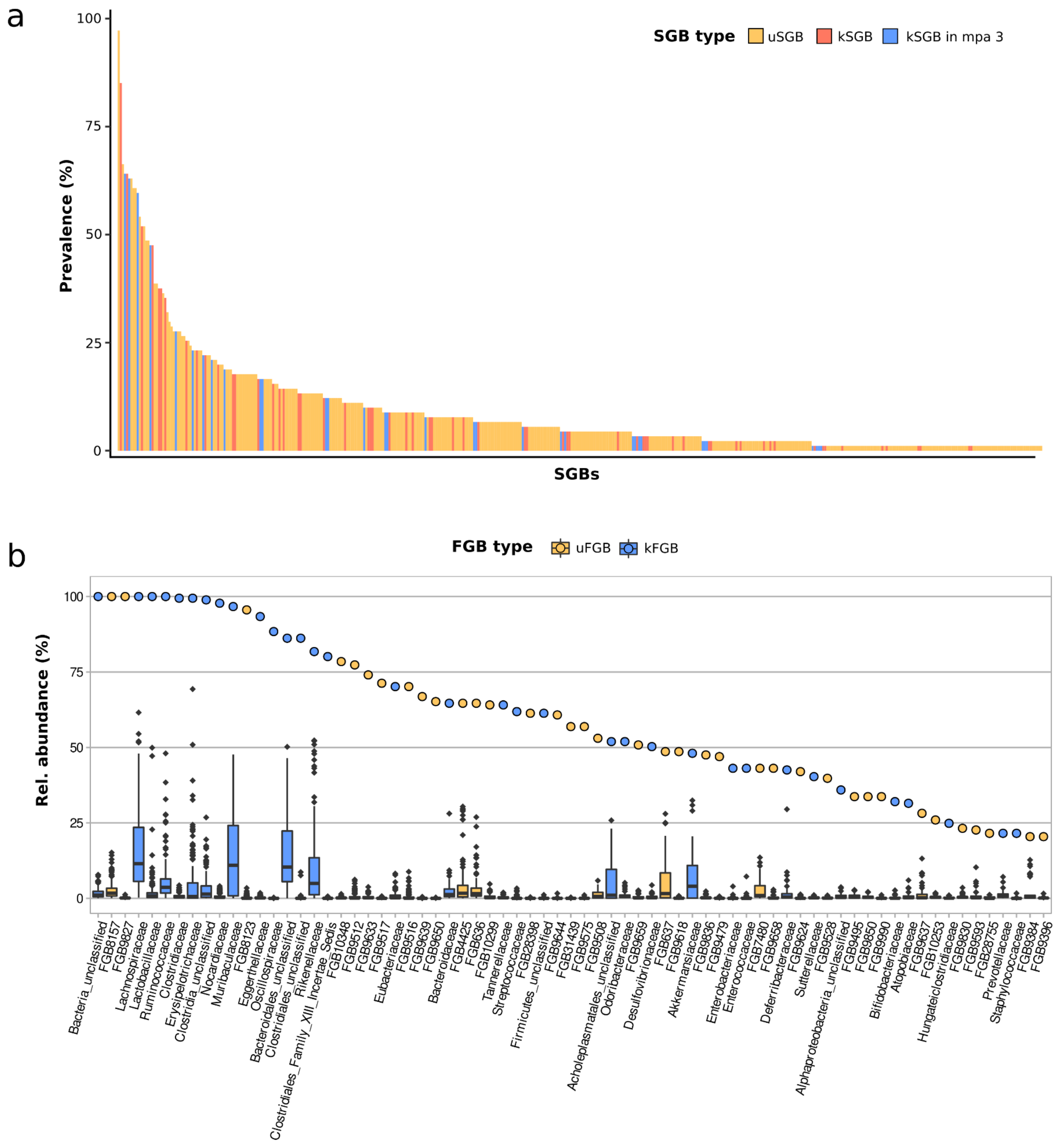


**Supplementary Figure 8. Expanded microbial diversity of the mice gut microbiome.** (**a**) The MetaPhlAn 4 genomic database incorporates 3,678 MAGs from the Xiao, L. et al 2015 study [61] which span 437 different SGBs. The proportion of kSGB and uSGB based on these MAGs follows a distribution similar to that based on the MetaPhlAn 4 taxonomic profiles (**Fig. 3a**). (**b**) Relative abundance and prevalence of the microbial families (FGBs) present in the mice gut. FGBs present in more than 20% of the samples are shown.

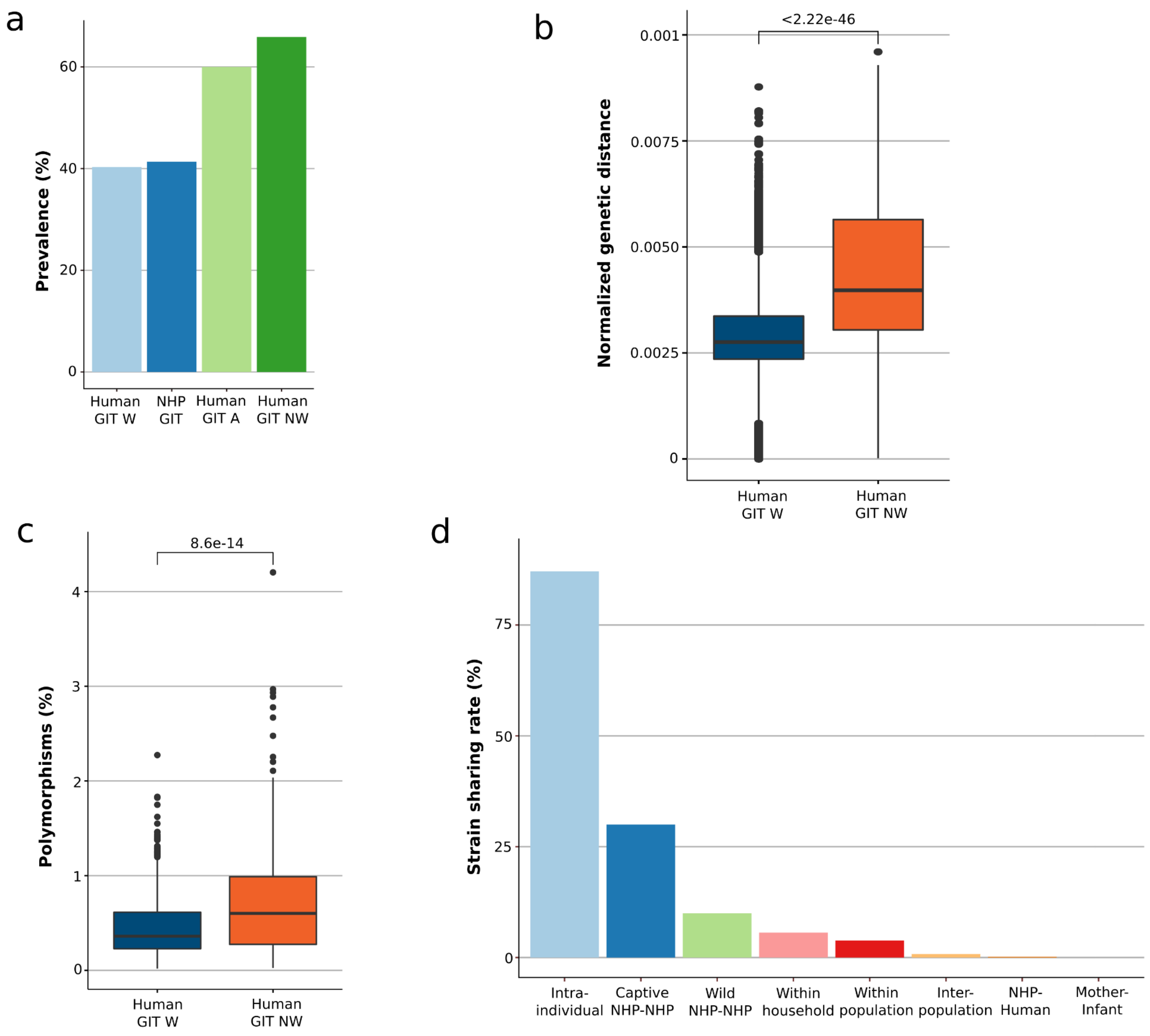


**Supplementary Figure 9. Phylogenetic analysis of *Lachnospiraceae* SGB4894.** (**a**) Prevalence of *Lachnospiraceae* SGB4894 across different hosts. For the modern human gut, only samples from healthy individuals were selected. (**b**) Intra-population normalized genetic distances differences between Westernized and non-Westernized populations (Mann-Whitney U test < 2.22e-46). (**c**) Polymorphic rates differences between Westernized and non-Westernized populations (Mann-Whitney U test = 8.6 e-14). (**d**) Strain sharing rates of *Lachnospiraceae* SGB4894. GIT=gastrointestinal tract, W=Westernized, NW=non-Westernized, A=ancient, NHP=non-human primates.

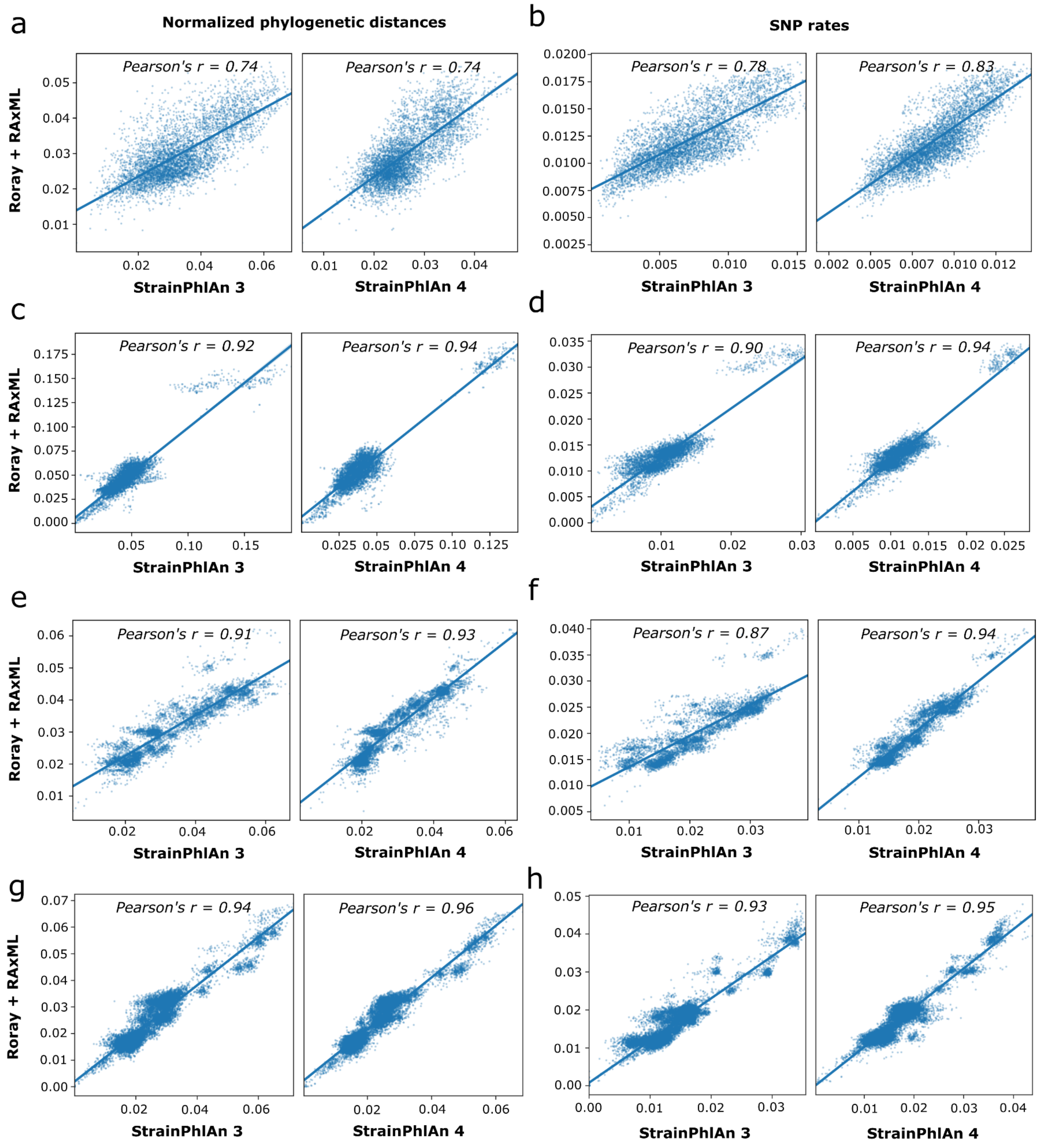


**Supplementary Figure 10. Comparison between StrainPhlAn 3 and 4 phylogenetic reconstruction.** Correlation of the normalized pairwise phylogenetic distances (**a,c,e,g**) and SNP rates (**b,d,f,h**) between the StrainPhlAn (x axis) and the Roary + RAxML (y axis) trees for (**a,b**) *Blautia wexlerae* (SGB4837), (**c,d**) *Bacteroides uniformis* (SGB1836), (**e,f**) *Eubacterium rectale* (SGB4933) and (**g,h**) *Lachnospiraceae* SGB4894.
